# Gene expression in the developing nemertean brain indicates convergent evolution of complex brains in Spiralia

**DOI:** 10.1101/2021.03.29.437382

**Authors:** Ludwik Gąsiorowski, Aina Børve, Irina A. Cherneva, Andrea Orús-Alcalde, Andreas Hejnol

## Abstract

**Background:** Nemertea is a clade of worm-like animals, which belongs to a larger animal group called Spiralia (together with e.g. annelids, flatworms and mollusks). Many of the nemertean species possess a complex central nervous system (CNS) with a prominent brain, and elaborated chemosensory and neuroglandular cerebral organs, which have been suggested as homologues to the annelid mushroom bodies. In order to understand the developmental and evolutionary origins of complex nemertean brain, we investigated details of neuroanatomy and gene expression in the brain and cerebral organs of the juveniles of nemertean *Lineus ruber*.

**Results:** In the hatched juveniles the CNS is already composed of all major elements present in the adults, including the brain (with dorsal and ventral lobes), paired longitudinal lateral nerve cords and an unpaired dorsal nerve cord. The TEM investigation of the juvenile cerebral organ revealed that the structure is already composed of several distinct cell types present also in the adults. We further investigated the expression of twelve transcription factors commonly used as brain and cell type markers in bilaterian brains, including genes specific for annelid mushroom bodies. The expression of the investigated genes in the brain is region-specific and divides the entire organ into several molecularly distinct areas, partially overlapping with the morphological compartments. Additionally, we detected expression of mushroom body specific genes in the developing cerebral organs.

**Conclusions:** At the moment of hatching, the juveniles of *L. ruber* already have a similar neuroarchitecture as adult worms, which suggests that further neural development is mostly related with increase in the size but not in complexity. Comparison in the gene expression between *L. ruber* and the annelid *Platynereis dumerilii* and other spiralians, indicates that the complex brains present in those two species evolved convergently by independent expansion of non-homologues regions of the simpler brain present in their common ancestor. The similarities in gene expression in mushroom bodies and cerebral organs might be a result of the convergent recruitment of the same genes into patterning of non-homologues organs or the results of more complicated evolutionary processes, in which conserved and novel cell types contribute to the non-homologues structures.

## Background

Nemertea is a clade of ca. 1300 described species of unsegmented worms, which predominantly occur in marine environments [1-3]. Phylogenetically, they belong to the large animal group called Spiralia (together with e.g. annelids, mollusks and flatforms) [4-12], however, despite recent progress in molecular phylogenetics, their exact position on the spiralian tree of life remains controversial [6-8, 10, 13].

Most nemerteans are active predators, which hunt for their invertebrate prey using a specialized eversible proboscis, a morphological apomorphy of the clade [1, 14-18]. This active lifestyle is accompanied by a relatively complex nervous system, composed of a large, multilobed brain (with two ventral and two dorsal lobes), a pair of lateral medullary nerve cords, extensive peripheral network and multiple specialized sensory organs [17-29]. Among the latter, the most conspicuous are the so-called cerebral (or cephalic) organs – paired structures of neurosecretory and either chemo- or mechanosensory function, located on the lateral sides of the head [17-23, 28, 30-33]. The exact arrangement of the cerebral organs varies between nemertean clades from relatively simple ciliated pits present in some Tubulaniformes, to the complex neuroglandular structures connected both directly to the brain and, through the convoluted ciliated canal, to the external environment in lineid heteronemerteans [17-23, 27, 28, 32, 33]. The phylogenetic analysis of morphological traits in nemerteans indicated that cerebral organs were already present in the last common nemertean ancestor [20]. However, it remains unclear, whether the cerebral organs represent an autapomorphy of nemerteans or homologs to some organs present in other spiralians such as ciliated pits of flatworms [30, 34] or mushroom bodies of annelids [19, 35, 36].

In the present study, we describe the detailed morphology of the nervous system and gene expression in the brain and cerebral organs of the juveniles of *Lineus ruber* (Müller, 1774), a directly developing lineid heteronemertean. *L. ruber* has been studied in past for both adult morphology [20, 22-26, 29-31] and some aspects of its development [29, 37, 38], including the molecular patterning of anterior-posterior axis, germ layers and lateral nerve cords [39, 40]. Comparison of our data with the existing morphological descriptions of the adult nervous system in *L. ruber* [20, 22-26, 29-31] and other closely related species, allows a better understanding of the ontogeny of the complex nemertean nervous system. Additionally, juxtaposition of gene expression profiles in the developing brain of *L. ruber* with that of other Spiralia [39, 41-48] can pinpoint similarities and differences in the molecular patterning of the spiralian brains in general, which in turn can inform evolution of the complex nemertean brain. Moreover, by comparing gene expression in cerebral organs of *L. ruber* and mushroom bodies of a comprehensively studied annelid *P. dumerilii* [49], we can provide new data to test the homology hypothesis of the cerebral organs of nemerteans and mushroom bodies of annelids.

## Results

### *Morphology of the nervous system in the juvenile* L. ruber

The investigated juveniles of *L. ruber* were freshly hatched from the egg mass, 42 days after oviposition [40]. We visualized the nervous system of the juveniles by applying antibody staining against tyrosinated tubulin, FMRF-amide and serotonin (5-HT), as well as Sytox green nuclear staining and fluorescent *in situ* mRNA hybridization of the choline acetyltransferase (*ChAT*), a genetic marker of the cholinergic neurons [50].

42 days old juveniles have already all major components of the nervous system (Figs. 1 and 2), which is composed of: 1) central nervous system (CNS) with brain, two lateral nerve cords (LNCs) connected by a postpharyngeal and posterior commissures and a single dorsal nerve cord (DNC); 2) stomatogastric nervous system (SNS), especially well developed in the pharyngeal region; 3) innervation of the proboscis; 4) network of fine peripheral nerves; 5) a pair of large cerebral organs; and 6) other sensory structures such as frontal organs and frontal sensory nerves.

**Figure 1.**
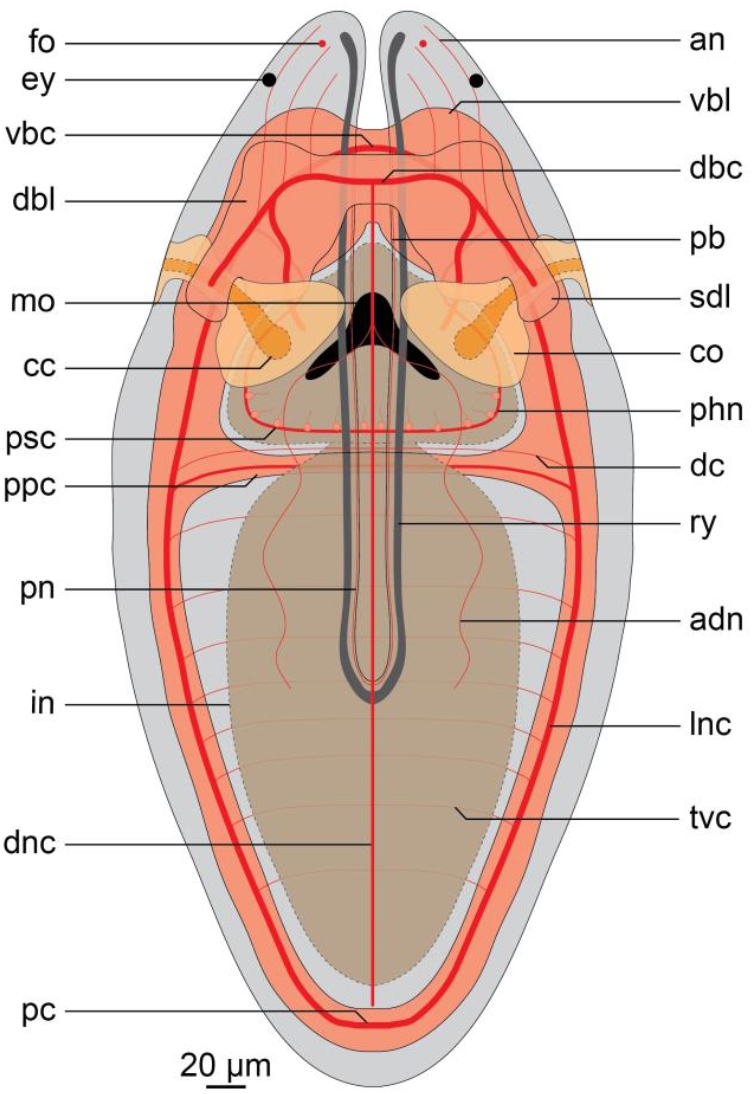
Schematic drawing of the nervous system in 42 days old juveniles of *Lineus ruber*. Anterior is to the top. Abbreviations: *adn* accessory dorsal nerve, *an* anterior nerve, *cc* ciliated canal, *co* cerebral organ, *dbc* dorsal brain commissure, *dbl* dorsal brain lobe, *dc* dorsal commissure, *dnc* dorsal nerve cord, *ey* eye, *fo* frontal organ, *in* intestine, *lnc* lateral nerve cord, *mo* mouth opening, *pb* proboscis, *pc* posterior commissure, *phn* pharyngeal nerve, *pn* proboscis nerve, *ppc* postpharyngeal commissure, *psc* pharyngeal sensory cell, *ry* rhynchocoel, *sdl* superior branch of the dorsal lobe, *tvc* transverse ventral commissure, *vbc* ventral brain commissure, *vbl* ventral brain lobe.

**Figure 2.**
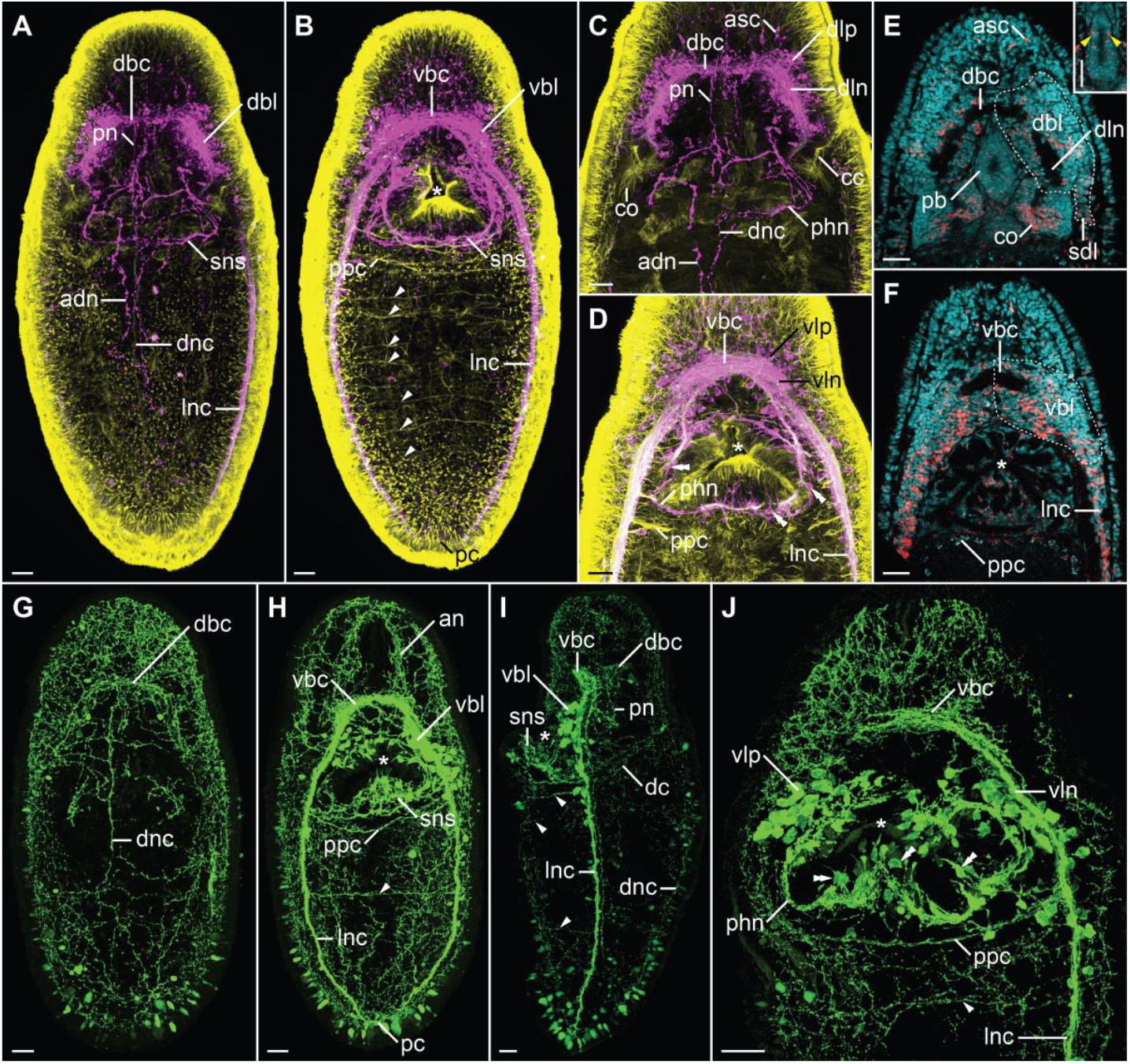
Morphology of the nervous system in 42 days old juveniles of *L. ruber* visualized with CLSM and antibody staining against tyrosinated tubulin (*yellow*, panels **A**–**D**), FMRF-amide (*magenta*, panels **A**– **D**) and serotonin (*green*, panels **G**–**J**) as well as Sytox green nuclear staining (*cyan*, panels **E, F**) and *in situ* hybridization with probe against choline acetyltransferase (*red*, panels **E, F**). Entire animal in dorso-ventral projection with a focus on dorsal (**A, G**) and ventral (**B, H**) structures; anterior part of the animal in dorso-ventral projection with a focus on dorsal (**C, E**) and ventral (**D, F, J**) structures, *inset* in panel **E** shows *ChAT* expression in the proboscis (*yellow arrowheads*); **I** lateral projection of the entire animal. Anterior is to the top on all panels. Scale bars 20 μm. Abbreviations: *adn* accessory dorsal nerve, *an* anterior nerve, *asc* anterior sensory cell, *cc* ciliated canal, *co* cerebral organ, *dbc* dorsal brain commissure, *dbl* dorsal brain lobe, *dc* dorsal commissure, *dln* dorsal lobe neuropile, *dlp* dorsal lobe perikaryon, *dnc* dorsal nerve cord, *lnc* lateral nerve cord, *pb* proboscis, *pc* posterior commissure, *phn* pharyngeal nerve, *pn* proboscis nerve, *ppc* postpharyngeal commissure, *sdl* superior branch of the dorsal lobe, *sns* stomatogastric nervous system, *vbc* ventral brain commissure, *vbl* ventral brain lobe, *vln* ventral lobe neuropile, *vlp* vetral lobe perikaryon. *White arrowheads* indicate transverse ventral commissures, *double white arrowheads* pharyngeal sensory cells and *asterisks* the mouth opening.

The brain is located anteriorly and is divided into four lobes: two ventral (*vbl*, Figs. 1 and 2B, F, H, I) and two dorsal ones (*dbl*, Figs. 1 and 2A, E). Each lobe is composed of the internal neuropile and the external layer of perikarya (Fig. 2C–F, J). Anteriorly both dorsal and ventral lobes are connected by dorsal (*dbc* Figs. 1 and 2A, C, E, G, I) and ventral (*vbc* Figs. 1 and 2B, D, F, H–J) brain commissures, respectively. Thus, the brain neuropile forms a ring around rhynchocoel and proboscis (Fig. 1). Posteriorly, each dorsal brain lobe is further divided into an inferior and a superior branch. The former connects directly to the cerebral organ (see below), while the latter ends blindly on the dorsal side of the animal (Figs. 1 and 2E). The neuropiles of the ventral lobes posteriorly give rise to the LNCs (Fig. 2D, H, J). FMRF-amide-like immunoreactive (FLIR) perikarya and *ChAT*^+^ cells have been observed in both dorsal and ventral brain lobes (Fig. 2A – F), while serotonin-like immunoreactive (SLIR) perikarya are present only in the ventral lobes (Fig. 2H–J). Both dorsal and ventral commissures and neuropiles of all brain lobes are composed of FLIR, SLIR and tyrosinated tubulin-like immunoreactive (TLIR) neurites (Fig. 2 A–D, G–J).

Three longitudinal nerve cords originate from the brain: a pair of thick LNCs (*lnc*, Figs. 1 and 2A, B, D, F, H–J) and a finer, unpaired DNC (*dnc*, Figs. 1, 2A, G, I). The LNCs are composed of an external layer of perikarya and an internal neuropile (and hence represent medullary nerve cords [51]). The neuropiles are densely packed with TLIR, SLIR and FLIR neurites (*lnc*, Fig. 2 A, B, D, H–J), while numerous *ChAT*^+^ neuronal cell bodies as well as more sparsely distributed FLIR and SLIR perikarya are mostly present in the anterior section of each LNC (Fig. 2B, D, F, I, J). The LNCs are connected behind the pharynx by a medullary postpharyngeal commissure (*ppc*, Figs. 2B, F, H, J), which is composed of TLIR and SLIR neurites as well as few SLIR and numerous *ChAT*^+^ perikarya (Fig. 2F and J). At the end of the animal body, both LNCs converge in a posterior commissure (*pc*, Figs. 1, 2B, H), which shows the same immunoreactivity patterns as neuropiles of LNCs. The DNC originates from the dorsal brain commissure. Compared to the LNCs, it is much finer and does not seem to be associated with any perikarya (Figs. 2A, G, I). It is composed of only a few TLIR and SLIR neurites, while anteriorly, a pair of fine FLIR dorsal accessory nerves branch out from it (*adn*, Figs. 1 and 2A). At the level of the pharynx, a fine, SLIR and TLIR dorsal commissure connects dorsal and lateral nerve cords (*dc*, Figs. 1 and 2I).

The SNS is composed of thick TLIR, FLIR and SLIR pharyngeal nerves, which originate from the ventral brain lobes and meander around the pharynx (*phn*, Figs. 1, 2C, D, J). Numerous sensory FLIR and SLIR cells are located along the pharyngeal nerves (*psc*, Fig. 1; *double arrowheads* Fig. 2D, J). Each of those cells has a basal connection to the pharyngeal nerve and an apical process pointing towards the pharyngeal lumen.

Some neural structures are also associated with the proboscis. Two longitudinal TLIR and FLIR nerves extend along the proboscis (*pn*, Figs. 1 and 2C), however their exact origin in the brain remains unclear. Scattered *ChAT*^+^ cells, of probably sensory function, are present in the epidermis of the proboscis (*yellow arrowheads*, inset in Fig. 2E).

The extensive network of peripheral nerves was detected, especially evident on the ventral side of the animal. It is composed of regular transverse ventral TLIR commissures (*tvc*, Fig. 1; arrowheads, Fig. 2B), some of which are additionally SLIR (*arrowheads*, Fig. 2H–J). A less regular network of SLIR intraepidermal neurites is present on both dorsal and ventral sides of the juvenile (Fig. 2 G–J).

A pair of conspicuous cerebral organs is located on the lateral sides of the head, just behind the brain (*co*, Figs. 1, 2C and E). More details of their morphology can be found in the following section. Other sensory structures, detected in addition to the cerebral organs, includes FLIR and *ChAT*^+^ anterior sensory cells (*asc*, Figs. 1 and 2 C, E), which likely contribute to the so-called frontal organs [19, 22, 23, 25], and numerous SLIR cephalic nerves extending anteriorly from the brain (*an*, Figs. 1 and 2H). Although 42 days old juveniles already possess rudiments of eyespots [40], we were not able to conclusively detect them in our investigation.

EdU staining in 60 days old juveniles showed that most of the brain cells at this later developmental stage are not mitotically active in contrast to the cells in other organs, such as proboscis, rhynchocoel or cerebral organs (Fig. 3A and B).

**Figure 3.**
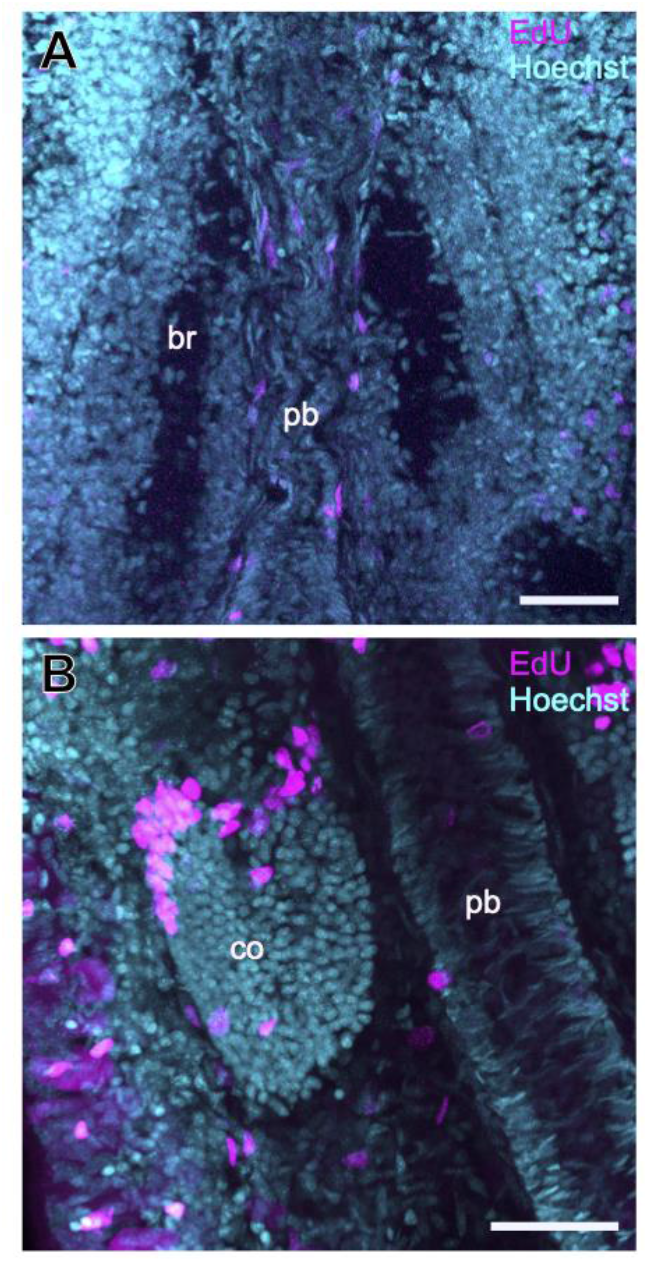
Proliferating cells in the head of 60 days old juveniles of *L. ruber* visualized by incorporation of EdU (*magenta*), counterstained with nuclear marker Hoechst (*cyan*). Dorso-ventral Z-projections of brain region (**A**) and cerebral organ (**B**), with anterior to the top. Scale bars 25 μm. Abbreviations: *br* brain, *co* cerebral organ, *pb* proboscis.

### Detailed morphology of the cerebral organs

Each cerebral organ is composed of two parts: a distal ciliated canal (*cc*, Figs. 1, 2C, 4B, C), which opens to the exterior on the side of the head (in the posterior part of the so called lateral cephalic slit), and a proximal neuroglandular portion (*co*, Figs. 1, 2C). The lumen of the ciliated canal is slightly curved in 42 days old juveniles, but the characteristic triple right-angle bends, present in the adult lineids [30-32] are not yet evident (*cc*, Fig. 2C). The ciliated canal connects the external environment with the neuroglandular part, which itself is firmly attached to the superior branch of the dorsal brain lobe (Fig. 1, 2E, and 4B, C). A thick TLIR and FLIR nerve of cerebral organ extends from the most posterior part of the dorsal lobe neuropile and penetrates the neuroglandular portion of the cerebral organ (*con*, Fig. 4C). We detected a few FLIR and much more numerous *ChAT*^+^ cells in the neuroglandular portion of the organ (*arrowhead*, Fig. 4C and *arrow*, Fig. 4B, respectively), while serotonin-like immunoreactivity was not detected (data not shown).

**Figure 4.**
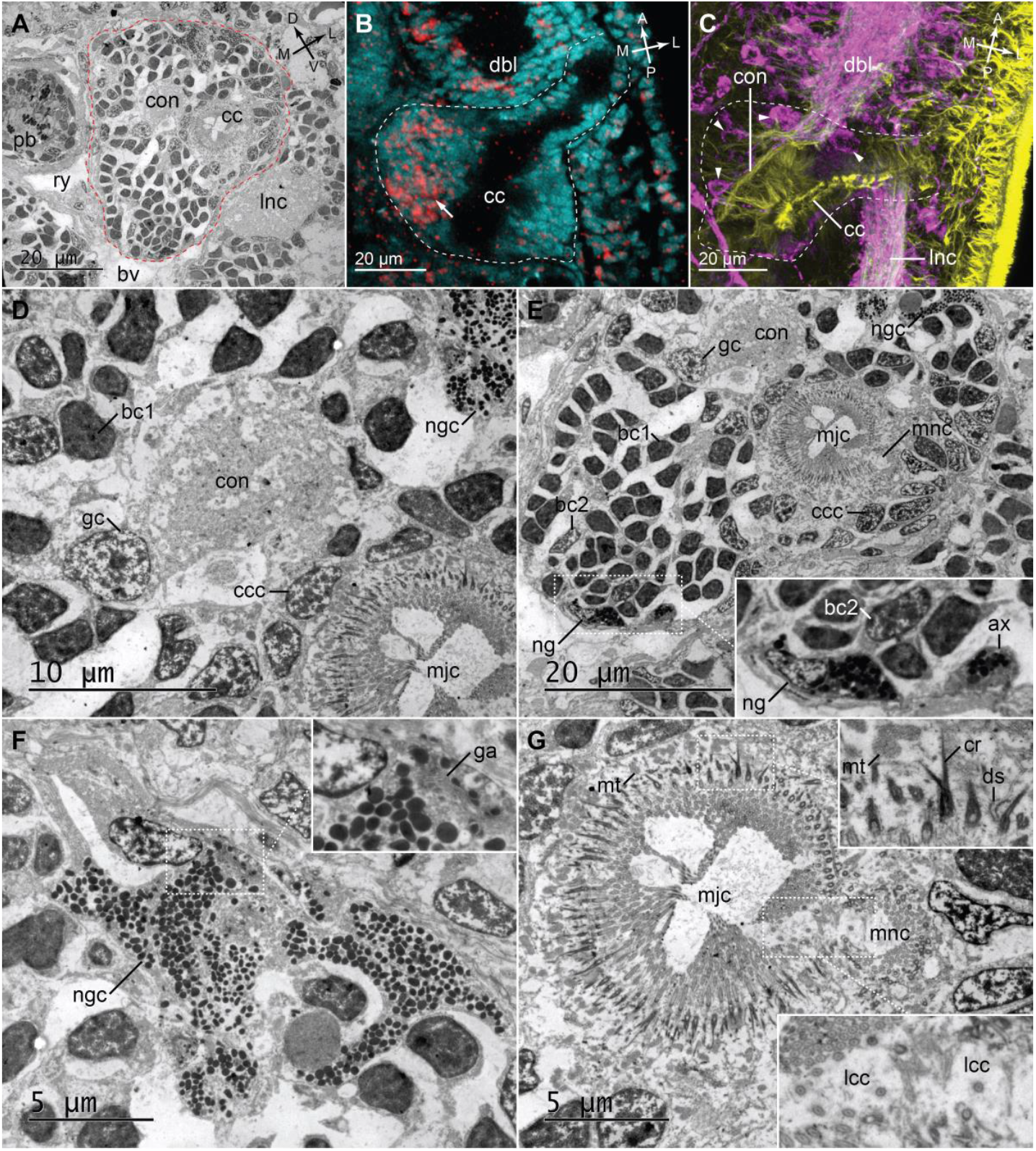
Detailed morphology of cerebral organs in juveniles of *L. ruber*. TEM micrographs of cerebral organs in 60 days old juvenile, showing cross section (**A**) and details of particular regions of the organ (**D**–**G**). Z-projections of cerebral organs in 42-days old juveniles visualized with Sytox green nuclear staining and *in situ* hybridization with probe against *ChAT* (*cyan* and *red*, respectively; **B**) and antibodies against FMRF-amide and tyrosinated tubulin (*magenta* and *yellow*, respectively; **C**). Cerebral organs are outlined in *red* (**A**) and *white* (**B, C**). Orientation inside the animal is indicated in the top-right corners in panels **A**–**C** (A, anterior; P, posterior; D, dorsal; V, ventral; M, median; L, lateral). Micrographs in panels **D**–**G**, show magnified areas of panel **A**. White outlined boxes on panels **E, F, G** indicates areas magnified in corresponding insets. Abbreviations: *ax* neuroglia axon, *bc1* bipolar cell type1, *bc2* bipolar cell type 2, *bv* blood vessel, *cc* ciliated canal, *ccc* ciliated canal cell, *con* nerve of cerebral organ, *cr* ciliary rootlet, *dbl* dorsal brain lobe, *ds* desmosome, *ga* Golgi apparatus, *gc* ganglion cell, *lcc* dilated cilia of lappet cell, *lnc* longitudinal nerve cord, *mjc* major ciliated canal, *mnc* minor ciliated canal, *mt* mitochondrium, *ng* neuroglia, *ngc* neuroglandular cell, *pb* proboscis, *ry* rhynchocoel. *White arrow* indicates ChAT^+^ cells in cerebral organ, *white arrowhead* FMRF-amide-like immunoreactivity in cerebral organ.

To gain further insight into the morphology of the cerebral organs, we supplemented the afore-mentioned confocal laser scanning microscopy (CLSM) based methods with ultrathin sectioning of resin-embedded specimens (60 days old juveniles) and TEM examination of the organ. That allowed us to describe the fine structure of the cerebral organ and ultrastructure of the particular cell types contributing to it. Since all detected cell types correspond directly to the ones described previously by Ling in his investigation of adult *L. ruber* [30], we adopted the terminology used therein.

We investigated cross-sections through the neuroglandular portion of the cephalic organ. The mass of the organ is located between the proboscis and the lateral nerve cords (Fig. 4A) and it is penetrated by both the cerebral organ nerve (*con*) and the ciliated canal (*cc*). The ciliated canal is divided into two parallel parts: a larger major ciliated canal (*mjc*) and a smaller minor ciliated canal (*mnc*) (Fig. 4G). Based on the ultrastructure, six distinct cell types can be distinguished in the sectioned area of the cerebral organ. The most numerous are type 1 bipolar cells (*bc1*), which constitute the majority of the cells in the neuroglandular mass (Fig. 4D, E). Their relatively small nuclei are roughly polygonal in cross-section and have dark nucleoplasm with the irregularly distributed chromatin (Fig. 4D). The very similar type 2 bipolar cells (*bc2*) are much less frequent (Fig. 4E). They have the same nuclear size and shape as well as chromatin arrangement as *bc1*, but their nucleoplasm is electron-translucent (Fig. 4E). A relatively few ganglion cells (*gc*) are present in the vicinity of the nerve of cerebral organ (Fig. 4D, E). Those cells have large nucleus that is almost circular in section and displays an electron-translucent nucleoplasm with nucleolus and irregularly distributed chromatin (Fig. 4D). On the dorsolateral side of the cerebral organ a single, large, irregularly shaped cell has been identified as neuroglandular cell (*ngc*, Fig. 4D–F). Its branching, spacious cytoplasm is filled with numerous electron-dense inclusions. Additionally, the Golgi apparatus was observed in the cytoplasm (*ga*; inset, Fig. 4F). A single neuroglial cell (*ng*) was observed on the opposite, ventro-median side of the organ (Fig. 4E). It is less voluminous than the neuroglandular cell, has a darker cytoplasm and more densely packed inclusions. A structure interpreted as a neuroglial axon is visible ca. 3 μm from the neuroglial cell body (*ax*; inset, Fig. 4E). The cells of the ciliated canal (*ccc*) represent the last cell type visible on the examined cross section (Fig. 4D, E). The apical surface of those cells is densely packed with cilia, which are equipped with asymmetrically bifurcating ciliary rootlets (*cr*; inset, Fig. 4G). Numerous mitochondria are present just below the ciliary rootlets, while the lateral sides of the cells are connected apically by desmosomes (*mt* and *ds*, respectively; inset, Fig. 4G). The cilia on the border of the major and the minor canals (*lcc*) are characteristically dilated and form a septum that divides both canals (inset, Fig. 4G). Those cilia indicate the presence of the seventh cell type, lappet cells, although the cells themselves could not be told apart from the other cells of the ciliated canal.

EdU staining of mitotically active cells in the 60 days old juveniles indicted intensive proliferation in cerebral organs, especially in its anterior region (Fig. 3B).

### Gene expression in the head

We investigated expression of 12 transcription factors (TFs), which have a role in CNS development of many bilaterians. Those genes include the conserved general brain markers (*otx, bf1*), genes involved in brain regional specification (*pax6, nk2*.*1, nk2*.*2, rx, otp*) and other neural genes, which are co-expressed in the annelid mushroom bodies (*dach, emx, arx, svp, tll*).

Expression of *otx* has been previously described for earlier developmental stages of *L. ruber*, in which the gene has a general anterior expression in the head [40]. In the 42 days old juveniles, which we investigated, the gene *otx* is predominantly expressed in the brain (Fig. 5A and B) and cerebral organs (Figs. 5A, 6B). In the brain, *otx* is broadly and uniformly expressed both in dorsal and ventral lobes (Fig. 5A and B). In the cerebral organs it is also widely expressed, both in the ciliated canal and neuroglandular part (Fig. 6B). A similar expression pattern of *otx* in the brain and cerebral organs has been also reported from developing juveniles of closely related *Lineus viridis* [52].

**Figure 5.**
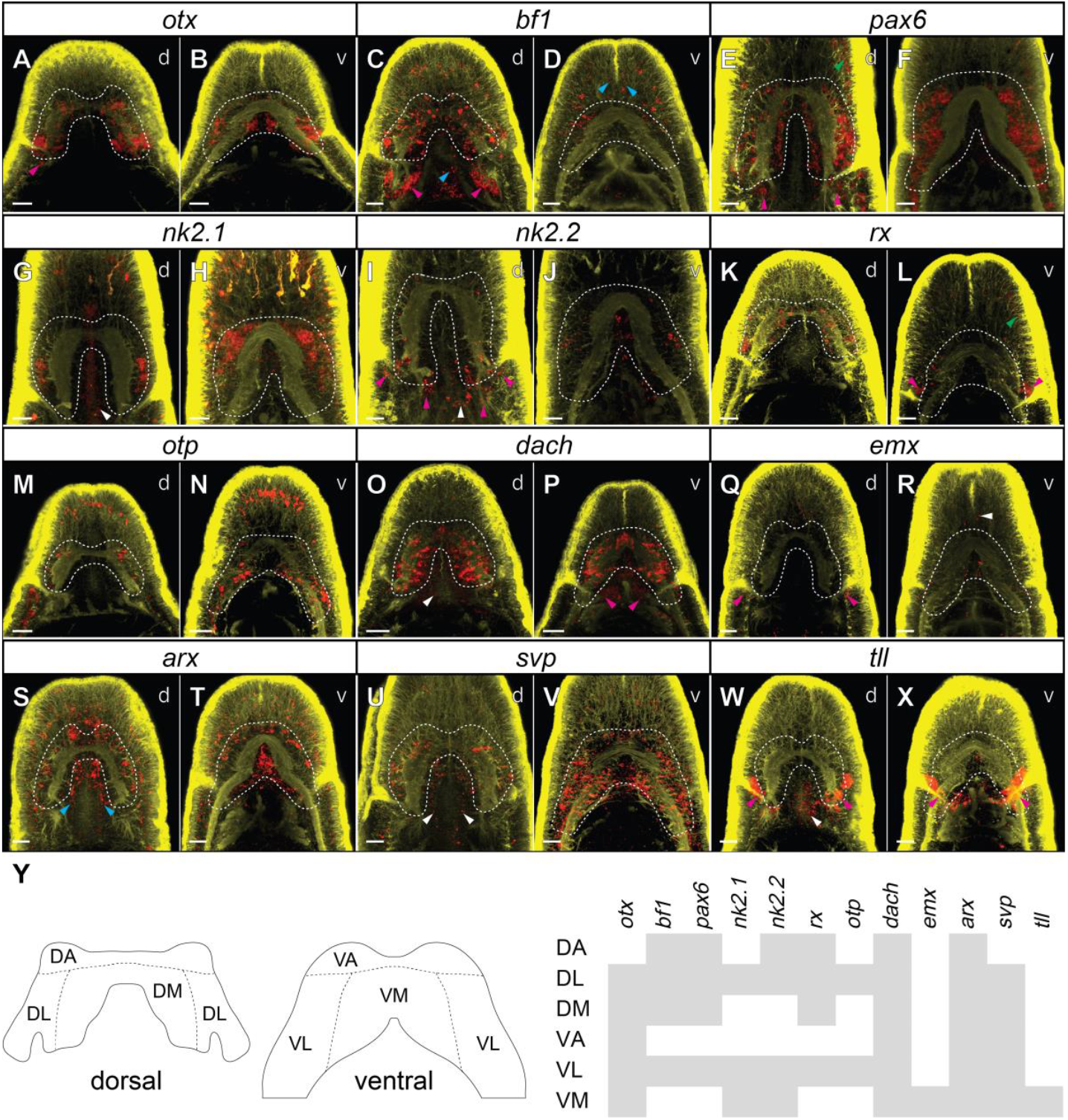
Expression of investigated transcription factors in the heads of 42 days old juveniles of *L. ruber*. **A**–**X** fluorescent *in situ* RNA hybridization, for each panel the name of the hybridized gene is shown in the white box above the micrographs. Fluorescent signal from RNA probes is in *red*, from antibody staining against tyrosinated tubulin in *yellow* and brain lobes are outlined in *white*. All animals are shown in dorso-ventral projection with anterior to the top; the letter in the top-right corner of each panel indicates whether focus is on dorsal (*d*) or ventral (*v*) structures. Detailed expression patterns are described in the text. *Magenta* arrowheads indicate expression in the cerebral organs, *blue* in the rhynchocoel, *green* in the lateral cephalic slits, *white* in the proboscis. Scale bars 20 μm. **Y** map of gene expression in the *L. ruber* brain. Grey bars indicate that gene is expressed in a particular brain region. Abbreviations: *DA* dorso-anterior brain domain, *DL* dorso-lateral brain domain, *DM* dorso-median brain domain, *VA* ventro-anterior brain domain, *VL* ventro-lateral brain domain, *VM* ventro-median brain domain.

**Figure 6.**
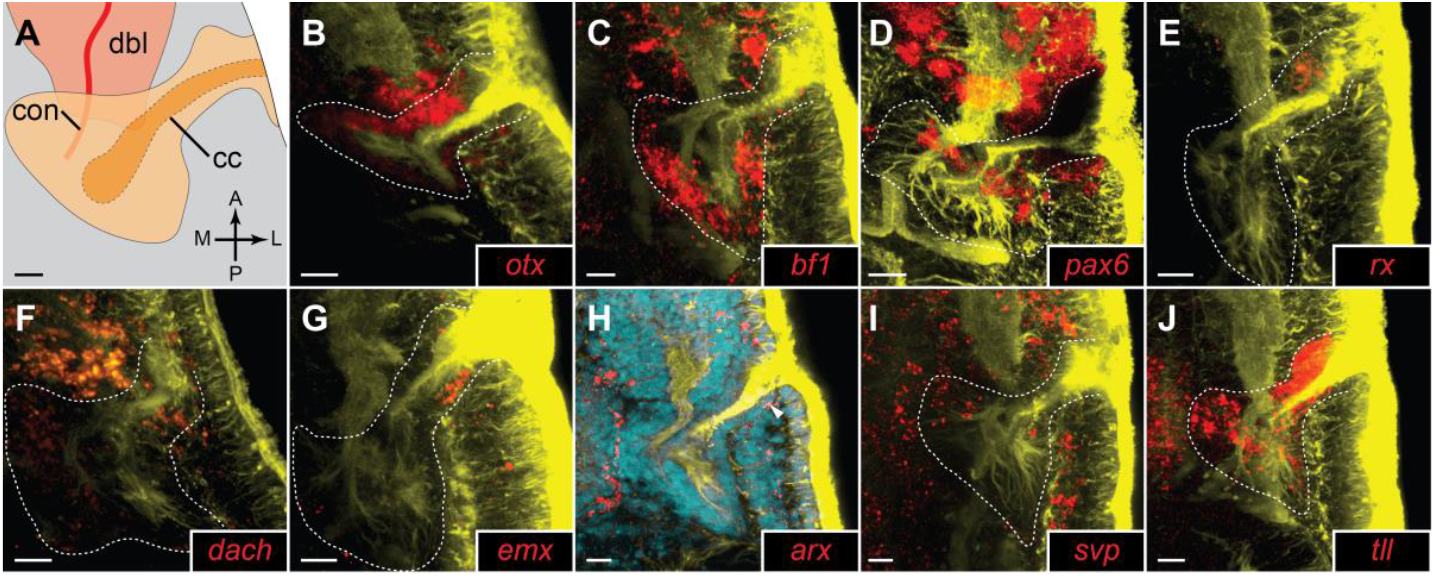
Details of gene expression in the cerebral organs of 42 days old juveniles of *L. ruber*. **A** schematic drawing of the cerebral organ and accompanying neural structures, orientation in the animal is indicated in bottom-right corner (A, anterior; P, posterior; M, median; L, lateral). Abbreviations: *cc* ciliated canal, *con* nerve of cerebral organ, *dbl* dorsal brain lobe. **B**–**J** fluorescent *in situ* RNA hybridization, for each panel the name of hybridized gene is provided in the bottom-right corner. Fluorescent signal from RNA probes is in *red*, from antibody staining against tyrosinated tubulin in *yellow* and from Sytox green nuclear staining in *cyan*; cerebral organs are outlined in *white*. The detailed expression patterns are described in the text. White arrowhead indicates *arx*^+^ cell at the posterior side of the ciliated canal opening. Scale bars 10 μm.

*bf1* is expressed in the brain, cerebral organs, scattered cells in the anterior epidermis and in the rhynchocoel (Figs. 5C and D, 6C). In the brain *bf1* is broadly expressed in the dorsal lobe (Fig. 5C), but in the ventral one it is only detectable in the lateral clusters of cells (Fig. 5D). The detected expression of *bf1* in the cerebral organ is very strong in the neuroglandular part, whereas we did not detect a signal in the ciliated canal (Fig. 6C).

Expression of *pax6, nk2*.*1* and *nk2*.*2* has been previously investigated in the juveniles *of L. ruber* in relation to the nerve cord patterning [39], however, the expression of those three genes in the brain was not described in the details that we provide here. In the head region, *pax6* is expressed in the brain, the epidermal cells of the lateral cephalic slits and in the cerebral organs (Figs. 5E and F, 6D). The gene is broadly expressed in the dorsal lobes (Fig. 5E), while in the ventral ones its expression is restricted to the lateral portions of the brain (Fig. 5F). In the cerebral organs the gene is expressed in the stripe of cells on the lateral side of the neuroglandular portion (Fig. 6D).

In the head region, *nk2*.*1* is expressed in the brain and proboscis (Fig. 5G and H). In the dorsal lobes the gene is expressed only in the small lateral clusters of cells (Fig. 5G), while on the ventral side the gene is broadly expressed both in the median and lateral domains (Fig. 5H). *nk2*.*1* is not expressed in the cerebral organs.

*nk2*.*2* is expressed in the brain, proboscis and cerebral organs (Fig. 5I and J). In the dorsal brain lobes, the gene is expressed in large clusters of posterior cells and in scattered anterior domains (Fig. 5I), whereas ventrally, it is expressed in median and lateral cell clusters (Fig 5J). Expression in the cerebral organs is detected in isolated domains of both ciliated canal and neuroglandular portion (Fig. 5I).

The gene *rx* is expressed in the brain, anterior sensory organs, epidermal cells of lateral cephalic slits and in the cerebral organs (Figs. 5K and L, 6E). Dorsally, the gene is expressed in isolated cells distributed relatively uniformly throughout the brain lobes (Fig. 5K). In the ventral lobes, *rx* is expressed only in a pair of postero-lateral cell clusters (Fig. 5L). In the cerebral organs, the gene is specifically expressed in the cluster of epidermal cells at the anterior side of the ciliated canal opening (Fig. 6E).

Expression of *otp* is detectable in the brain, LNCs, and numerous anterior sensory cells (Fig. 5M and N). In the dorsal lobes, the gene is expressed only in a relatively few lateral cells (Fig. 5M), while ventrally it is also predominantly expressed in the lateral cells of the brain lobes, but its expression was also detected in the more median cells contributing to the mouth innervation and anterior part of the LNC (Fig. 5N).

In the head region, the gene *dach* is expressed in the brain, cerebral organs, proboscis and few isolated anterior cells (Figs. 5O and P, 6F). The expression in the brain is rather uniform and transcripts of the gene were detected in all regions of both dorsal and ventral lobes (Fig. 5O and P). In the cerebral organs, the gene was detected in some of the cells of both the ciliated canals and the neuroglandular portion (Fig. 6F).

Expression of the gene *emx* was detected in the brain, cerebral organs, proboscis, and cells along anterior cephalic nerves (Figs. 5Q and R, 6G). In the brain the gene is expressed only in a few cells in the ventro-median domain (Fig. 5R). In the cerebral organs the gene transcripts were detected in the cells at the posterior side of the ciliated canal opening and in a single median cell in the neuroglandular part of the organ (Fig. 6G).

The TF *arx* has a broad expression in the anterior body of the juvenile *L. ruber*. It is expressed in the brain, rhynchocoel, epidermal cells, anterior sensory cells and in the cerebral organs (Figs. 5S and T, 6H). In both dorsal and ventral brain lobes, its expression was detected in numerous anterior, lateral and median cells (Figs. 5S and T). In contrast, the expression in the cerebral organs was restricted to a single cell at the posterior side of the ciliated canal opening (Fig. 6H).

The gene *svp* is also broadly expressed in anterior structures; its expression was detected in the brain, cerebral organs, LNCs, anterior sensory cells and proboscis (Figs. 5U and V, 6I). In the dorsal brain lobes, it is expressed in cells distributed through the lateral and median regions (Fig. 5U), while ventrally it is expressed uniformly in the entire ventral lobes (Fig. 5V). In the cerebral organs, expression of *svp* was detected in some anterior and lateral cells of the neuroglandular part (Fig. 6I).

Transcripts of the gene *tll* were detected in the brain, cerebral organs and proboscis (Figs. 5W and X, 6J). Expression in the brain was restricted just to a few cells posteriorly to the ventral commissure (Fig. 5X). Signal from the probes against *tll* was extremely strong in the cerebral organs (Fig. 5W and X) and was observed throughout the entire structure in cells of both the ciliated canal and the neuroglandular portion (Fig. 6J).

The brain of the juvenile *L. ruber* is divided by commissures and lobe neuropiles into eight regions: unpaired dorso-anterior, dorso-median, ventro-anterior and ventro-median regions as well as paired dorso-lateral and ventro-lateral areas (Fig. 5Y). Mapping of the above-described gene expression patterns onto those brain domains reveled that most of the regions express unique combination of the TFs (Fig. 5Y). The only brain regions which seem to express the same sets of TFs are dorsal and ventral lateral domains (Fig. 5Y).

### Gene co-expression during brain development

To further explore co-expression of some of the TFs in the brain, we performed double *in situ* hybridization of the selected brain patterning genes (*nk2*.*1, nk2*.*2, pax6* and *rx*). In addition to the investigation of 42 days old juveniles, we also examined co-expression of those genes in the earlier developmental stage, 25 days old early juveniles, in order to test whether the observed co-expression patterns are conserved throughout ontogenesis.

The CNS of 25 days old juveniles shows much simpler morphology when compared to the hatched juveniles (Fig. 7A). It is composed of LNCs, which merge anteriorly in the brain with two commissures – a thicker ventral and thinner dorsal – that form a ring shaped neuropile around the developing proboscis rudiment. At this developmental stage, the brain is not yet divided into the dorsal and ventral lobes and the cerebral organs are not fully formed, being mainly composed by the ciliated canal, that is not directly connected with the brain [40].

**Figure 7.**
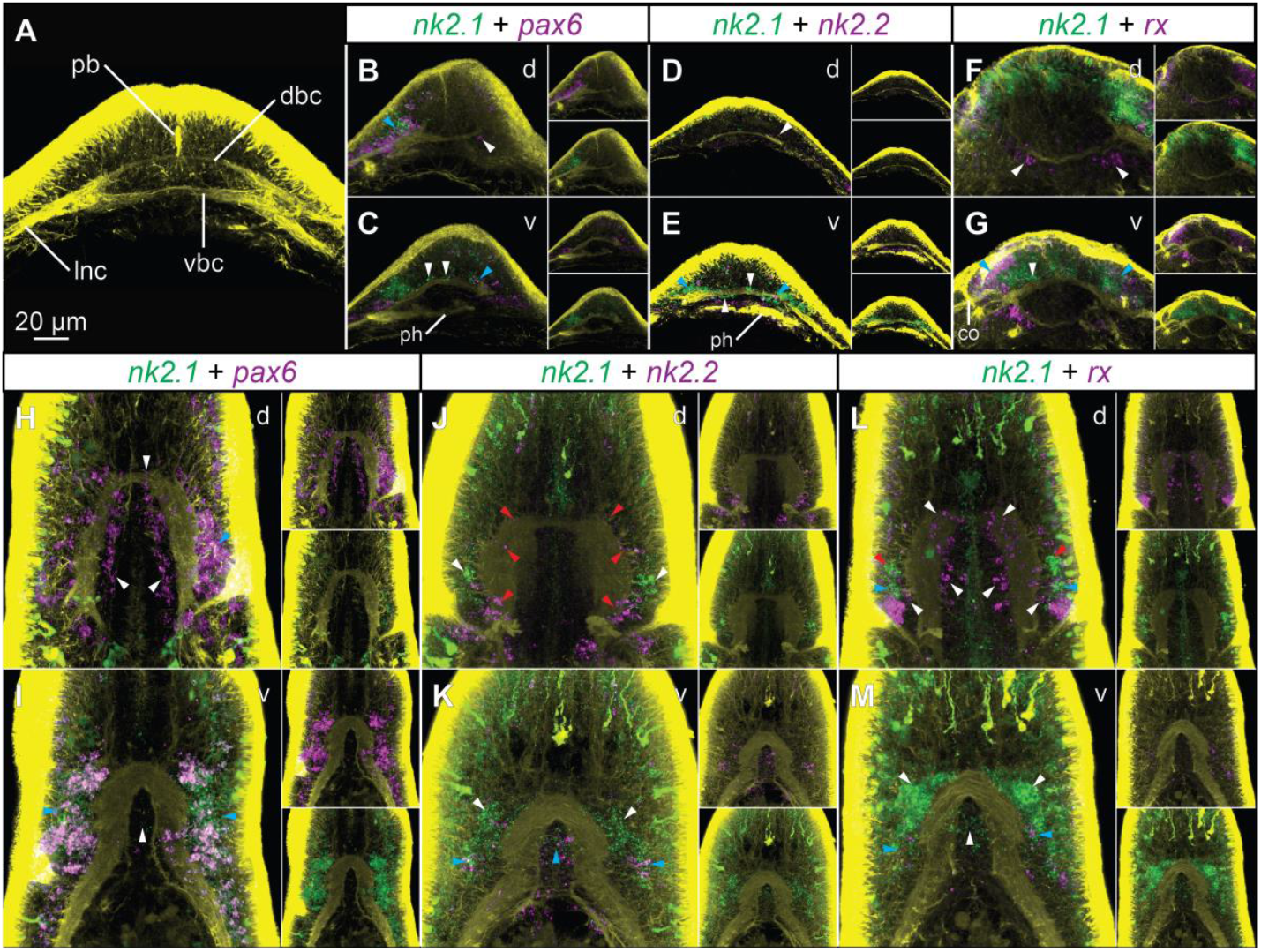
Co-expression of brain patterning genes in the developing brain of *L. ruber*. **A** morphology of the brain in 25 days old juveniles. **B**–**G** co-expression in the brain of 25-days old juveniles. **H**–**M** co-expression in the brain of 42-days old juveniles. For each panel the color-coded names of hybridized genes are shown in the white box above the micrographs. *White* and *red* arrowheads indicate exclusive expression of one of the hybridized genes, *blue* arrowheads indicate co-expression. All animals are shown in dorso-ventral projection with anterior to the top; the letter in the top-right corner of each panel indicates whether the focus is on dorsal (*d*) or ventral (*v*) structures. Micrographs on panels **B**–**M** are not to the scale. Abbreviations: *co* cerebral organ, *dbc* dorsal brain commissure, *lnc* lateral nerve cord, *pb* proboscis rudiment, *ph* pharynx.

In the brain of 25 days old juvenile, *nk2*.*1* is expressed along the ventral commissure and in the lateral parts of the brain (Fig. 7B, C, E, G). In its lateral domains the gene is co-expressed with *pax6* (blue arrowheads, Fig. 7B and C) and *rx* (blue arrowheads, Fig. 7G). Additionally, some of the lateral *nk2*.*1*^+^ cells also express *nk2*.*2* (blue arrowheads, Fig. 7E). The more median *nk2*.*1*^+^ cells that are associated with the ventral commissure are devoid of *pax6, nk2*.*2* and *rx* expression (white arrowheads, Fig. 7C, E, G). In addition to the expression in lateral domains, *pax6, nk2*.*2* and *rx* are also expressed in cells associated with the dorsal commissure, which do not co-express *nk2*.*1* (white arrowheads, Fig. 7B, D, F).

The analysis of gene co-expression in the 42 days old juveniles generally corroborates the expression map based on single gene hybridization, however it allows more detailed description of the brain molecular regionalization. In the dorsal brain *pax6* is broadly expressed in the lateral and median domains (white arrowheads, Fig. 7H) and only small clusters of lateral cells co-express *pax6* and *nk2*.*1* (blue arrowhead, Fig. 7H). In the ventral lobes, the lateral cells co-express *pax6* and *nk2*.*1* (blue arrowheads, Fig. 7I), while cells in the median domain express only *nk2*.*1* (white arrowhead, Fig. 7I). *nk2*.*1* and *nk2*.*2* are not co-expressed in the dorsal brain (Fig. 7J). *nk2*.*1* is expressed in the most lateral cells of the dorsal brain (white arrowheads, Fig. 7J), while *nk2*.*2* is expressed in the large, more posterior domains and in scattered cells in the anterior brain region (red arrowheads, Fig. 7J). In the ventral brain, both genes are co-expressed in the postero-lateral and median domains (blue arrowheads, Fig. 7K), however *nk2*.*1* has much broader ventral expression with many *nk2*.*1*^+^ cells devoid of *nk2*.*2* expression (white arrowheads, Fig. 7K). *rx* is expressed in scattered anterior, median and lateral cells in the dorsal brain, which do not co-express *nk2*.*1* (white arrowheads, Fig. 7L). In the lateral parts of the brain some cells co-express *rx* and *nk2*.*1* (blue arrowheads, Fig. 7L), while some *nk2*.*1*^+^ cells do not express *rx* (red arrowheads, Fig. 7L). In the ventral brain the antero-lateral and median *nk2*.*1*^+^ cells do not express *rx* (white arrowheads, Fig. 7M), while small clusters of postero-lateral cells co-express both genes (blue arrowheads, Fig. 7M).

On the whole, comparison of gene co-expression between 25- and 42-days old juveniles shows that the general molecular patterning of the developing brain is retained throughout development. The ventro-median region expresses *nk2*.*1* but not *pax6* nor *rx*. The lateral brain includes cells co-expressing *nk2*.*1* with *pax6, nk2*.*2* and *rx*, while dorsal brain is mainly composed of *pax6, nk2*.*2* and *rx* positive cells which do not co-express *nk2*.*1*. The differences between both life stages are primarily associated with the more complex architecture of the brain in 42 days old juveniles, which requires a more intricate developmental control, nevertheless the most general gene expression patterns are conserved.

## Discussion

### Comparison of juvenile and adult morphology

Nervous system has been investigated in great detail in adult *Lineus ruber* [20, 22-26, 29-31] and *Lineus viridis* [19, 20, 24, 25], a morphologically similar species that belongs to the same species complex [53, 54]. Comparison between the juvenile and adult worms reveals that all major nervous structures described in the adults are already present in the 42 days old juveniles, indicating that at this stage the general neuroarchitecture is already fully formed and that further development is mostly related with increase in the size but not morphological complexity. The same pattern is observed in number and diversity of cell types contributing to the cerebral organs. There are, however, some minor differences in immunoreactivity patterns between both life stages. For instance, SLIR perikarya have been reported in the dorsal brain ganglia of adult *L. ruber* [23], while we observed immunoreactivity against serotonin only in the ventral brain ganglia of the juveniles (Fig. 2I). This indicates, that even though the general morphology of the brain is already established at the moment of hatching, the following growth of the brain is not purely quantitative, but also new cell types are added in some brain regions during further development. Moreover, staining of mitotically active cells showed that in 60 days old juveniles cell proliferation in the brain is lower than in the other organs, while the cells of the cerebral organs are still intensively dividing (Fig. 3), indicating allometric growth of the CNS.

The major postpharyngeal commissure, which ventrally connects the lateral nerve cords, is the only juvenile neural structure which does not correspond directly to any of the elements of the adult nervous system of *L. ruber* [20, 22, 29] or, to our best knowledge, of any other nemertean, which nervous system has been studied thus far [e.g. 17-19, 20, 21, 28, 55-58]. In adult nemerteans, the lateral nerve cords are connected by numerous delicate ventral commissures, that are composed just of bundles of neurites and are considered as part of the peripheral nervous system. Conversely, the postpharyngeal commissure described in this study is associated with few SLIR and numerous *ChAT*^+^ perikarya and has typical medullary arrangement, markedly different from the remaining ventral commissures (Figs. 1 and 2). There are two possibilities to explain this discrepancy in morphology of both stages: either the commissure degenerates during ontogeny or, due to the allometric growth, becomes much less prominent in later developmental stages and was overlooked in previous investigations.

Nevertheless, the observation of the postpharyngeal ventral commissure in a nemertean is interesting since similar structures are present in numerous annelids (e.g. the first commissure connecting ventral nerve cords [59-64]), as well as in all major clades of gastrotrichs [65-67] and gnathiferans [68-71]. Therefore, the distribution of this character on the phylogenetic tree raises the possibility that the ventral postpharyngeal commissure connecting the major nerve cords might represent a plesiomorphic spiralian trait retained in some form in numerous clades.

### Expression of brain patterning genes in Spiralia

Molecular patterning of the brain has been investigated in relatively many spiralians, representing diverse clades with broad spectrum of morphological complexity of their brains (Tab. 1). Among those species, the best studied is the annelid *Platynereis dumerilii*, which possesses a relatively complex brain with multiple morphologically, functionally and developmentally distinct regions [42, 49, 72-74]. One of the important characteristics of gene expression patterns during the development of the *P. dumerilii* brain is regional restriction of *nk2*.*1* expression to the ventro-median region and *pax6* expression in the lateral domains (including eyes and mushroom bodies), with only the minimal overlap of expression of both genes (Fig. 8A; [42, 49]). This expression pattern resembles the one observed in vertebrates [75, 76] and has been proposed as an ancestral bilaterian trait [42]. Although a comparable expression of those two genes is also witnessed in some Spiralia (Tab. 1), including other annelids [43, 47, 48], rotifers [39] and brachiopods [39, 46, 77-79], we did not retrieve a similar pattern in neither 25-nor 42-days old juveniles of *L. ruber* (Figs. 7B, C, H, I and 8B). *nk2*.*1* is indeed mostly expressed in the ventral domain (Figs. 5H, 8B), however, it is broadly co-expressed with *pax6* in the ventral lobes and in the small dorso-lateral domains (Figs. 7H, J, 8B); while *pax6* shows expression not only in the lateral domains but is generally broadly expressed throughout the entire brain (including the dorso-median domain), with the only exception of the small ventro-median region (Figs. 5E, F, 8B). A very similar expression of *nk2*.*1* and *pax6* has been observed in planarians, where *nk2*.*1* is expressed mostly in the ventral portion of the brain [44, 45], while one of the *pax6* paralogs, *pax6A*, is broadly expressed in the brain tissue [44, 80]. A further parallel between planarians and *Lineus* is associated with a seemingly diminished role of *pax6* in eye formation: *pax6* is not expressed during eye development neither in *L. ruber* (this study) nor in *L. viridis* [81] (although it seems to have a role in eye regeneration in *L. sanguisues* [81]), while in flatworms eye regeneration has been demonstrated to be *pax6* independent [80]. The role of *pax6* in eye patterning is otherwise highly conserved among bilaterians [e.g. 82, 83, 84]. Due to the unstable position of Nemertea on the spiralian phylogeny [e.g. 6-8, 10], it is currently impossible to determine whether those similarities between platyhelminths and nemerteans are due to the convergent evolution, a common evolutionary innovation or retention of ancestral plesiomorphic conditions in both lineages.

**Table 1.**
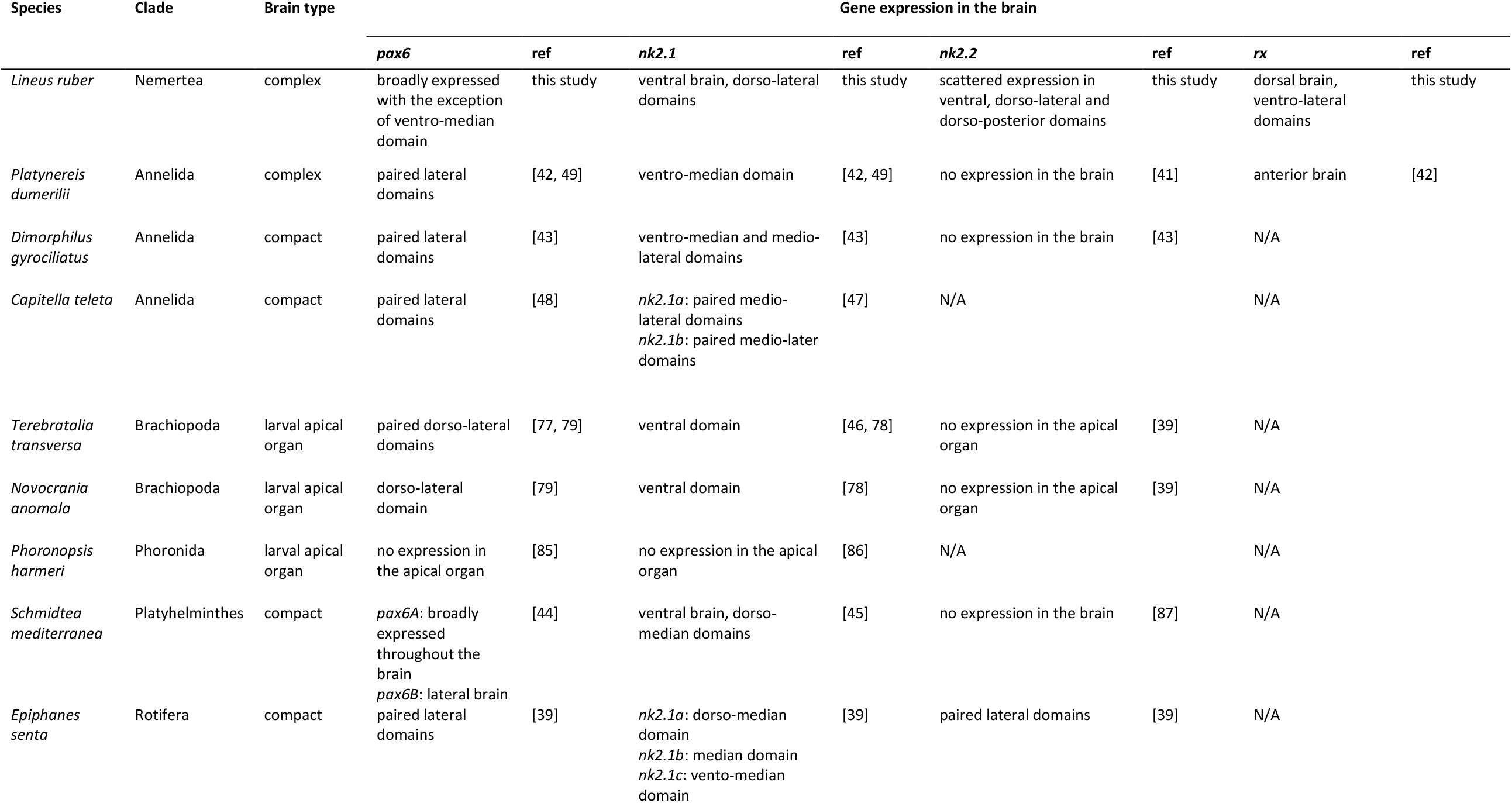
Expression of the selected genes in the spiralian brains.

**Figure 8.**
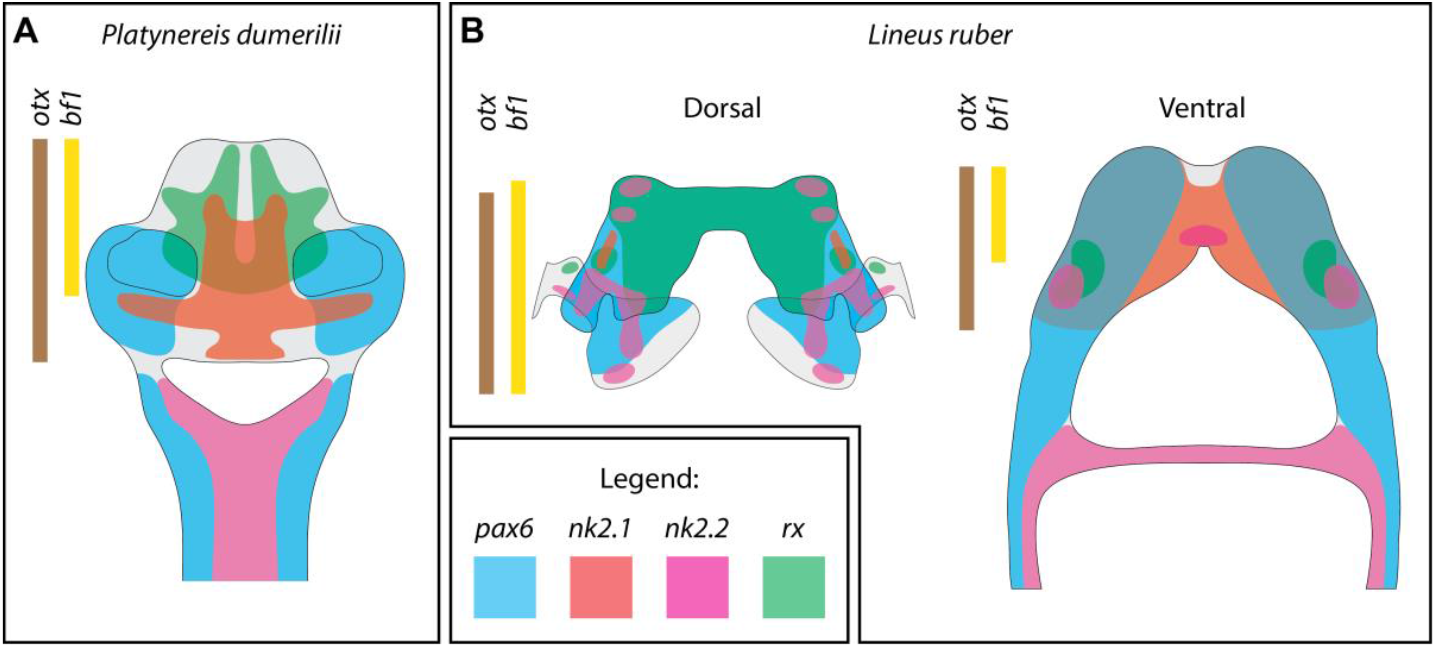
Comparison of gene expression in the CNS of (**A**) annelid *Platynereis dumerilii* (based on results from [41, 42, 49]) and (**B**) nemertean *Lineus ruber* (based on current study and [39]).

Another important differences in expression of brain patterning genes between *L. ruber* and other Spiralia includes the expression of *nk2*.*2* within numerous brain domains of *L. ruber* (while the gene lacks brain expression not only in annelids [41, 43], but also in brachiopods [39] and flatworms [87]) as well as broad expression of *rx* in the dorsal lobes of the nemertean brain (*versus* their more rostral expression in *P. dumerilii* [42, 49]). Altogether this comparison shows that complex brains of nemerteans, and especially their dorsal lobes, show little resemblance in the molecular patterning to the complex brains of *P. dumerilii* (Fig. 8), which in turns seem to share more molecular similarities with simpler brains of other annelids and apical organs of brachiopod larvae (Tab. 1). This observation, in concert with morphological data [21, 64, 88], indicates that complex brains of nemerteans and errant annelids evolved convergently, due to e.g. similar selective pressure associated with predatory/active life style [89]. We propose that the increase in the brain size and complexity in those two lineages was achieved by independent expansions of non-homologous regions of simpler brains present in their respective ancestors.

Some of the investigated nemertean brain patterning genes are also expressed in the proboscis (*nk2*.*1*, nk2.2, *dach, svp, tll*) and rhynchocoel (*bf1, arx*), two morphological apomorphies of Nemertea [15, 16]. Taking into account that the proboscis is a highly innervated structure [this study; also 15-20, 22, 23, 25, 26, 29, 57, 58, 90], the neuronal genes in the proboscis might be expressed in the developing neuronal network of the organ. Comparable results were obtained by body region-specific transcriptomics of the nemertean *Notospermus geniculatus*, in which expression of some of the neuronal markers (e.g. *elav, syt12*) was also detected in the proboscis [91]. Expression of *arx* and *bf1* in the rhynchocoel, a coelom derived structure [16, 92], seems more peculiar, since those genes have a generally conserved neuroectodermal expression in Bilateria [46, 93-97]. However, *arx* is also expressed in clade-specific morphological structures of brachiopods (in chaetal sacs and protegulum forming epithelium [98, 99]), annelids (in chaetal sacs [100]) and mollusks (in radula formative tissue [101]). Therefore, our data just further expand the list of potential co-options of *arx* into patterning of spiralian evolutionary novelties.

### Are mushroom bodies and cerebral organs derived from the same ancestral organs?

In numerous annelid brains, morphologically distinct structures, referred to as mushroom bodies, are present, which have allegedly chemosensory and cognitive functions [35, 36, 49, 73, 102-107]. There is an ongoing discussion on whether those structures are part of the ancestral annelid body plan or whether they evolved more recently in one of the annelid subclades [35, 73, 108]. However, their phylogenetic distribution (especially the lack of comparable structures in Palaeoannelida and Sedentaria [64, 88, 109]) favors the latter option [64, 88, 108, 110].

Nevertheless, morphologically similar structures are also present in Panarthropoda [36, 105, 111-114], which lead some authors to the idea that mushroom bodies-like structures were already present in the common protostome ancestor [36, 49, 73, 105]. Although similarities in molecular patterning of annelid mushroom bodies and vertebrate pallium led to the assumption that both structures originated from the same sensory and associative brain center of hypothetical ancestral bilaterians [49], such homology statements, based on observation of only two phylogenetically distant clades, are always at the best case highly tentative [89, 115].

Cerebral organs of nemerteans, in contrast to the annelid mushroom bodies, can be unequivocally reconstructed as present in the last common nemertean ancestor [20, 22, 28]. However, it remains unresolved whether they are nemertean evolutionary novelty or rather homologs of the mushroom bodies of annelids [19, 35, 36] or the lateral ciliated pits present in catenulids and macrostomids [30, 34, 116, 117], the two earliest sequentially branching platyhelminth clades [118]. Similarities between the mushroom bodies of annelids and the cerebral organs of nemerteans are rather superficial: the former are integral parts of the brain and are not connected to the external realm, while the latter are always contacting ambient environment and, especially in Hoplonemertea, might be spatially separated from the CNS [17, 28, 32, 33]. On the other hand, the function, general morphology, connectivity and fine structure of cerebral organs of nemerteans and ciliated pits of flatworms bear a strong resemblance [30, 34, 116, 117], making their homology much more likely. Taking into account the arrangement of the cerebral organs in various nemertean clades, the “ciliated pit” organization seems to represent an ancestral character state also in nemerteans [19-21]. If one accepts that the cerebral organs of nemerteans and ciliated pits of catenulids and macrostomids are homologues [34], then, depending on the phylogenetic position of nemerteans, there are two possible scenarios of their evolution: 1) If nemerteans are sister group to platyhelminths (Parenchymia hypothesis [7, 119]), then the ciliated pits-like structures represent a synapomorphy of Parenchymia. 2) On the other hand, if nemerteans are closer to annelids than flatworms [5, 6, 8, 10], then the presence of ciliated pits might represent a plesiomorphic condition, present also in the annelid ancestor.

In the face of the above-discussed concerns about the homology of mushroom bodies and cerebral organs, we were surprised to find that cells constituting the cerebral organs express the same set of transcription factors as mushroom bodies of annelids (with both structures being additionally free of *nk2*.1 expression). Although all nine of the annelid mushroom body markers, which expression we tested, were expressed in the cerebral organs of *L. ruber*, they were not co-expressed uniformly throughout the entire structure. Some genes (*otx, bf1, dach* and *tll*) were expressed in all regions of the organ, while others were restricted only to some cells in the neuroglandular portion (*pax6, emx, svp*) or the ciliated canal (*rx, emx, arx*). The complicated landscape of TFs expression in *L. ruber* correlates well with the fact, that the cerebral organs of 60 days old juveniles are already composed of numerous diverse cell types, including neurons, glia cells, glandular cells and ciliated epidermal cells (Fig. 4) as well as still dividing, possibly not fully differentiated, cells (Fig. 3). Unfortunately, with the resolution of our data, we were not able to pinpoint co-expression of particular TFs with specific cell types contributing to the organ. In *P. dumerilii* these TFs are also not expressed uniformly in the entire mushroom body and show regionalized expression [49], however, their regionalization does not simply correspond to the one observed in the cerebral organs of *L. ruber*. For example, *otx* and *tll* are expressed only in the subset of neurons constituting mushroom body, while expression of *pax6, arx* and *svp* is detected in most of the cells forming the organ [49]. Therefore, even though the same set of genes is expressed in both types of organs, their exact co-expression in particular cell types is probably divergent and the apparent similarities in gene expression profiles between both organs might be more superficial than they appear on the first sight.

A further problem with the interpretation of the gene expression patterns in the cerebral organs is related to the fact that, both in annelid and in nemertean, it remains unknown whether those TFs interact in the same gene regulatory network (GRN) or whether they are independently expressed in different, unrelated cell types. If they are part of the same GRN, then co-option of the ancestral regulatory program into patterning of non-homologues structures might explain the observed similarities. If indeed the ciliated pits-like structures, homologues to the cerebral organs of nemerteans, were present in the annelid ancestor (see above) it is possible to envision a recruitment of the established genetic control of those organs into the patterning of chemoreceptive portion of the brain in the ancestral errant annelid. On the other hand, if the genes are not part of the same GRN and instead act independently in particular cell types (which is supported by non-corresponding, region-specific expression of particular TFs in mushroom bodies and cerebral organs) a more complicated mechanism might account for the observed similarities. For instance, some of the cell types present in both organs might be homologues and derived from the common ancestor, but the organs containing those cell types are convergent and include other, unrelated and lineage-specific cell types. This could happen due to the reduction of the ciliated duct and the secretory cells and further integration of the neural part of the ancestral ciliated pits with the CNS in annelids. A solid phylogenetic position of Nemertea, analysis of function and interactions of the studied TFs as well as additional gene expression data from catenulids, macrostomids and Palaeoannelida are needed to ascertain on any of those evolutionary scenarios.

## Conclusions

In this study, we investigated the morphology and gene expression in the developing CNS of the nemertean *Lineus ruber*. At the moment of hatching, juveniles of *L. ruber* have already all major components of the adult nervous system, which indicates that further development is mostly related with increase in the size but not morphological complexity. This likeness corelates well with a similar predatory lifestyle of both juveniles and adults [40]. Comparison of gene expression in the brain of *L. ruber* and the annelid *P. dumerilii* [41, 42, 49] indicates that complex brains, observed in those two animal species, evolved convergently by independent expansion of non-homologues regions of simpler ancestral brains. Such scenario corresponds with the similar conclusions drawn by comparative morphology [21, 64, 88]. In contrast to the discrepancies in gene expression in the brains, we observed that the same set of transcription factors, which is expressed in the mushroom bodies of *P. dumerilii* [49] is also expressed in the cerebral organs of *L. ruber*. These similarities might be a result of convergent recruitment of the same GRN into patterning of non-homologue organs or indicators of the homology of some cell types contributing to mushroom bodies and cerebral organs that could evolve from the cell type present in the lateral chemosensory ciliated pits of the hypothetical spiralian ancestor. Further studies on the cell-type level and functional interactions of the studied TFs are needed to fully resolve the level of homology, or convergence, between mushroom bodies and cerebral organs.

## Methods

### Animal collection and morphological investigation

Adult specimens of *Lineus ruber* were collected near Bergen, Norway (Fanafjord; GPS coordinates: 60.251845N, 5.320947E). The animals had dark red coloration with wide pigment-free areas in the terminal part of the head. Animals were kept in the laboratory in filtered seawater at 14°C with a daytime cycle: 13 hours of sunshine and 11 hours of darkness. Collection of egg masses and desired developmental stages, animal fixation as well as antibody, nuclear and EdU stainings followed the already established protocols [40].

Specimens for TEM investigation were fixed in 4% PFA in PBS, rinsed in the same buffer, postfixed in 1% OsO4 diluted in PBS for 120 min at 4°C, rinsed again and dehydrated in graded ethanol/acetone series. The samples were embedded in Epon 812 resin (Sigma Aldrich) and cut to semi- and ultrathin sections with a diamond knife (Diatome Histo Jumbo) using ultramicrotome Leica EM UC6. The ultrathin cross sections of cerebral organ were placed on formvar-covered (Fluka) single slot copper grids and stained with 1% uranyl acetate and lead citrate.

### Gene expression analysis

Coding sequences for analyzed genes were identified in the transcriptome of *L. ruber* with the reciprocal TBLASTN search using orthologous protein sequences from *P. dumerilii*. Sequence of all of the newly identified genes were translated into protein sequences and aligned with reference sequences from other animals (Table S1). The alignments were trimmed either manually or with TrimAl software [120] and analyzed with FastTree v2.1 [121] in order to assess orthology of the analyzed genes (Figs. S1–5). All newly obtained sequences were submitted to GenBank (Accession numbers MW720144–MW720151).

Fragments of genes were amplified from cDNA library using specific primer pairs, cloned into pGEM-T Easy vectors (Promega, USA) and then transformed into competent *Escherichia coli* cells for amplification. Plasmid DNA was isolated and sequenced in both forward and reverse directions using T7 and SP6 primers to assure that the desirable genes were cloned. The antisense probes were transcribed from linearized DNA and labeled either with digoxigenin (for hybridization of single mRNA) or with dinitrophenol (for detection of second gene in double *in situ* hybridization). Whole mount *in situ* hybridization followed the same procedure as described for *L. ruber* juveniles in other studies [39, 40].

### Imaging and image processing

Samples for confocal laser scanning microscopy (antibody staining and *in situ* hybridization) were mounted in Murray’s clear and scanned in either Leica SP5 or Olympus FV3000 CLSM. Z-stacks of confocal scans were projected into 2D images in IMARIS 9.1.2. TEM microphotographs were obtained with Gatan ES500W camera mounted on transmission electron microscope Jeol JEM-1011. Both CLSM images and TEM micrographs were assembled in Adobe Illustrator CS6 into final figures. All the schematic drawings were done with Adobe Illustrator CS6.

## Acknowledgements

We are grateful to all present and former members of the Comparative Developmental Biology Group, University of Bergen, who helped with the collection and culturing of *Lineus viridis*. We also would like to thank Naëlle Barabé, who cloned and prepared probe against *dach* gene. All TEM studies were carried out at the Shared Research Facility “Electron microscopy in life sciences” at Moscow State University.

## Authors’ contributions

LG conducted gene search and orthology assessments, cloned genes, performed *in situ* hybridization, arranged figures and drafted the manuscript. AB performed antibody staining, searched and cloned genes and performed *in situ* hybridization. IAC prepared, examined and photographed samples for TEM. AOA searched and cloned genes, performed antibody and EdU stainings. AH designed and coordinated the study and contributed to the writing. All authors read, accepted and approved the final version of the manuscript.

---

**Figure S1.**
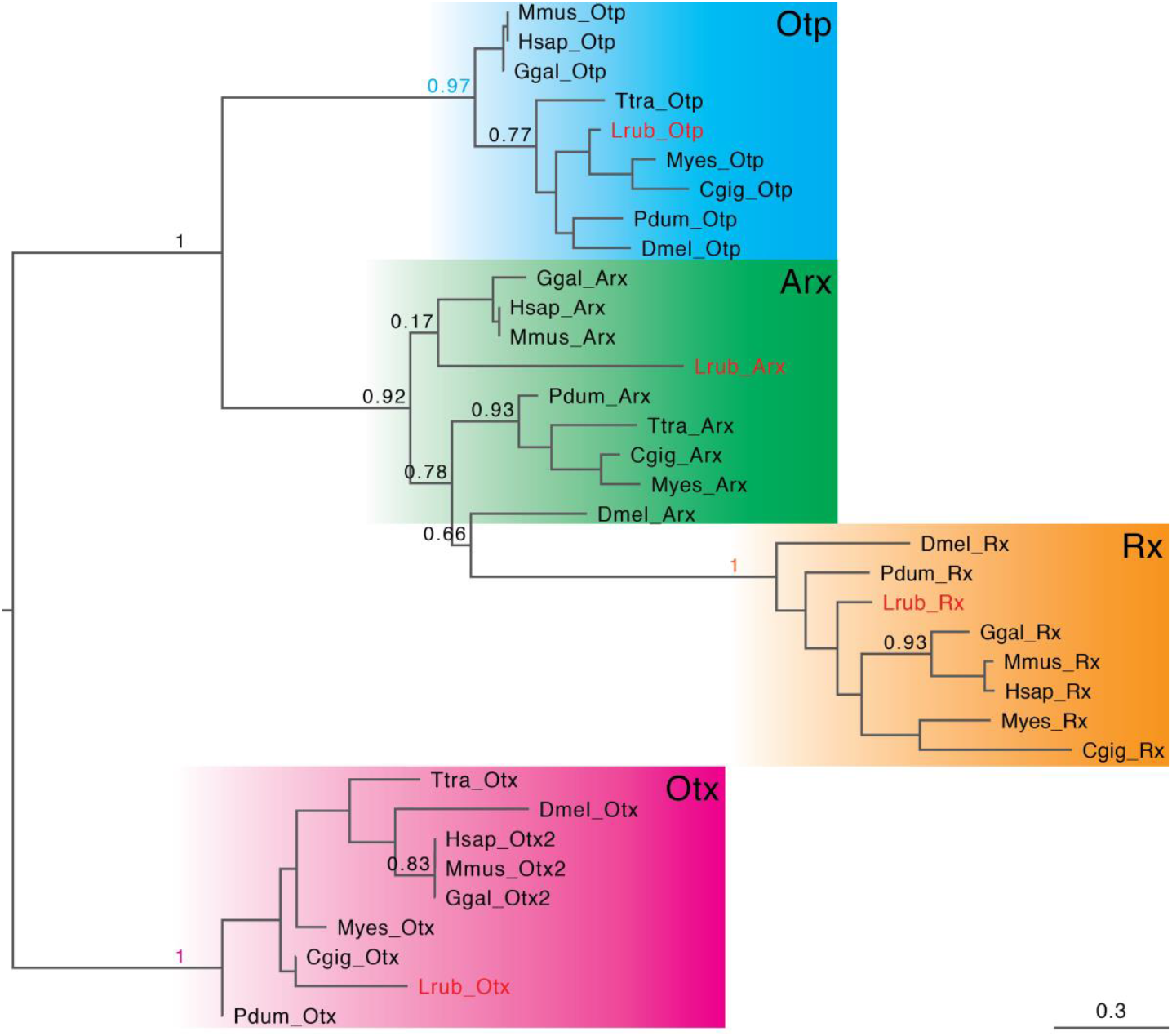
Phylogenetic analysis of PRD-class homeobox transcription factors. SH-like support values are shown for the important nodes. Scale bar on the lower right corner shows amino acid substitution rate per site. Sequences from *L. ruber* are marked in red. For abbreviation and source of other sequences see table S1.

**Figure S2.**
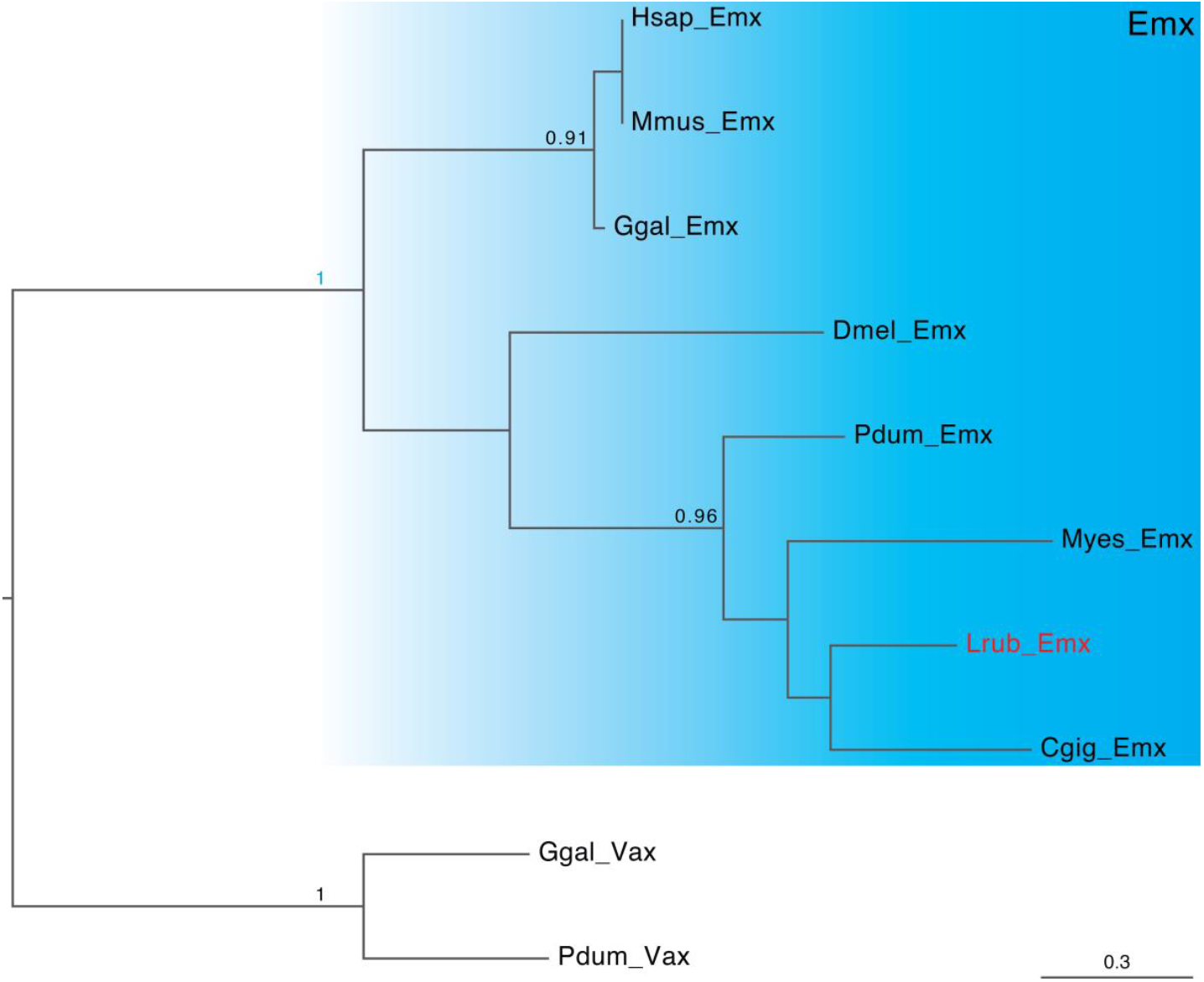
Phylogenetic analysis of Emx sequences. SH-like support values are shown for the important nodes. Scale bar on the lower right corner shows amino acid substitution rate per site. Sequence from *L. ruber* is marked in red. For abbreviation and source of other sequences see table S1.

**Figure S3.**
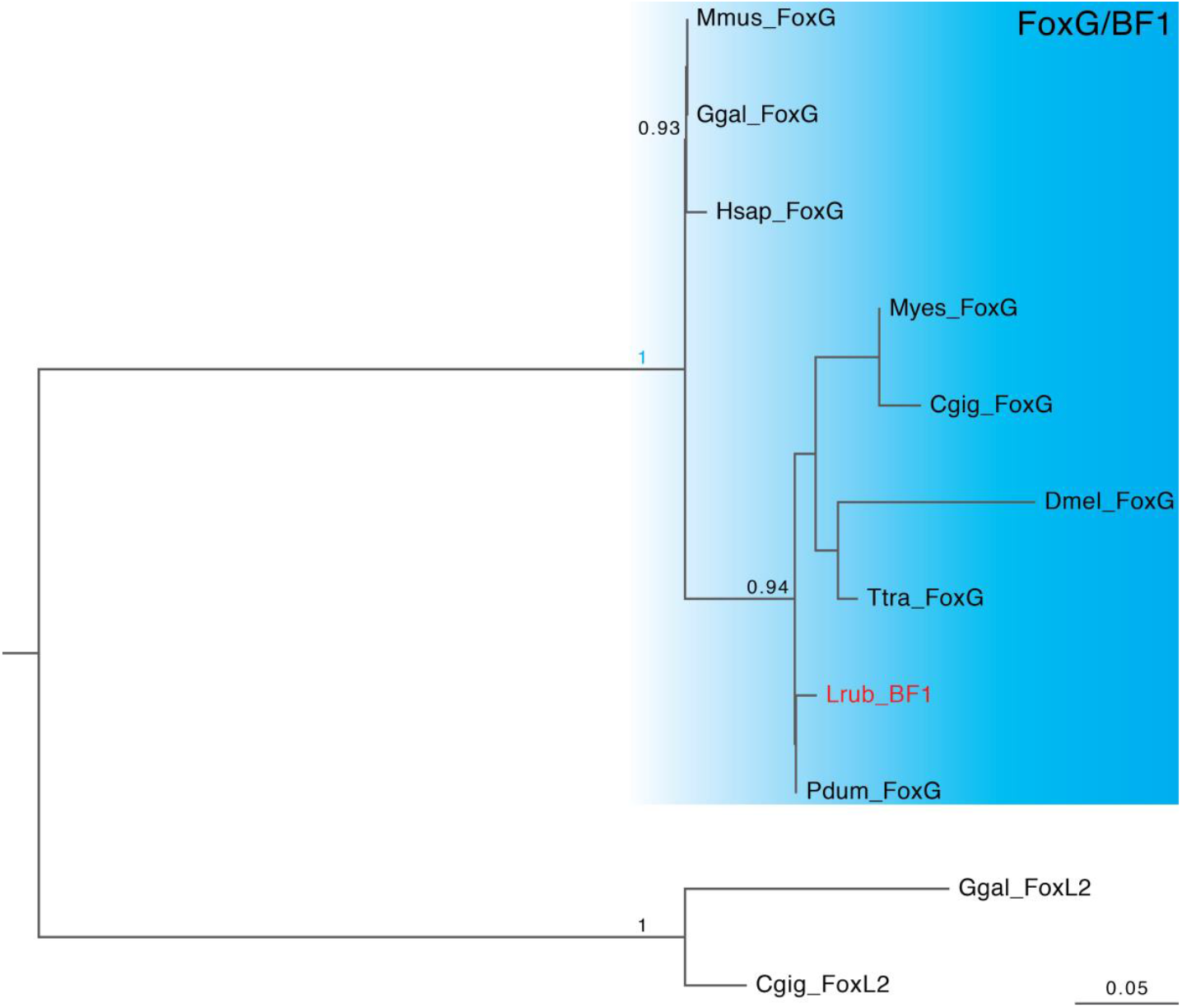
Phylogenetic analysis of Fox sequences. SH-like support values are shown for the important nodes. Scale bar on the lower right corner shows amino acid substitution rate per site. Sequence from *L. ruber* is marked in red. For abbreviation and source of other sequences see table S1.

**Figure S4.**
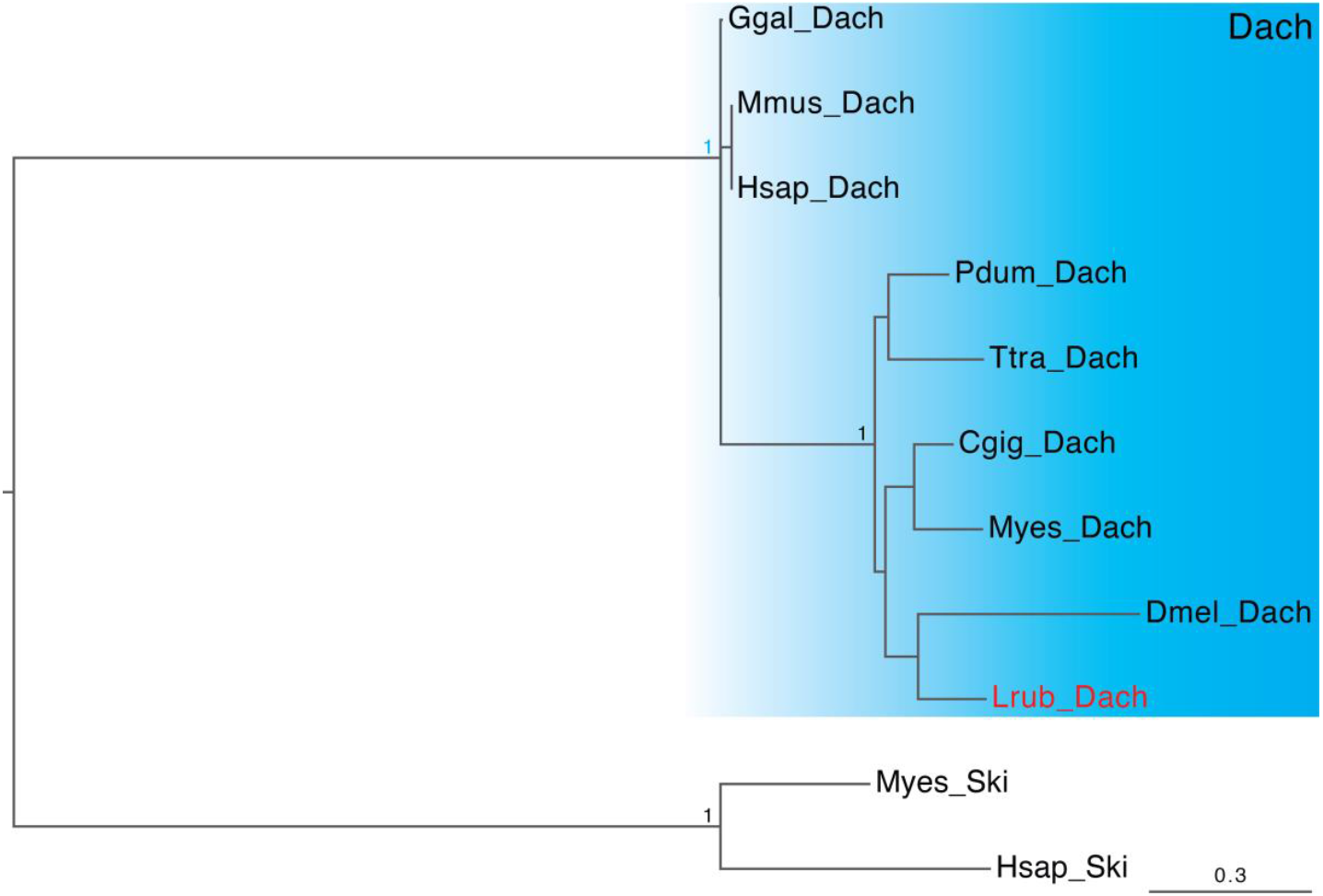
Phylogenetic analysis of Dach sequences. SH-like support values are shown for the important nodes. Scale bar on the lower right corner shows amino acid substitution rate per site. Sequence from *L. ruber* is marked in red. For abbreviation and source of other sequences see table S1.

**Figure S5.**
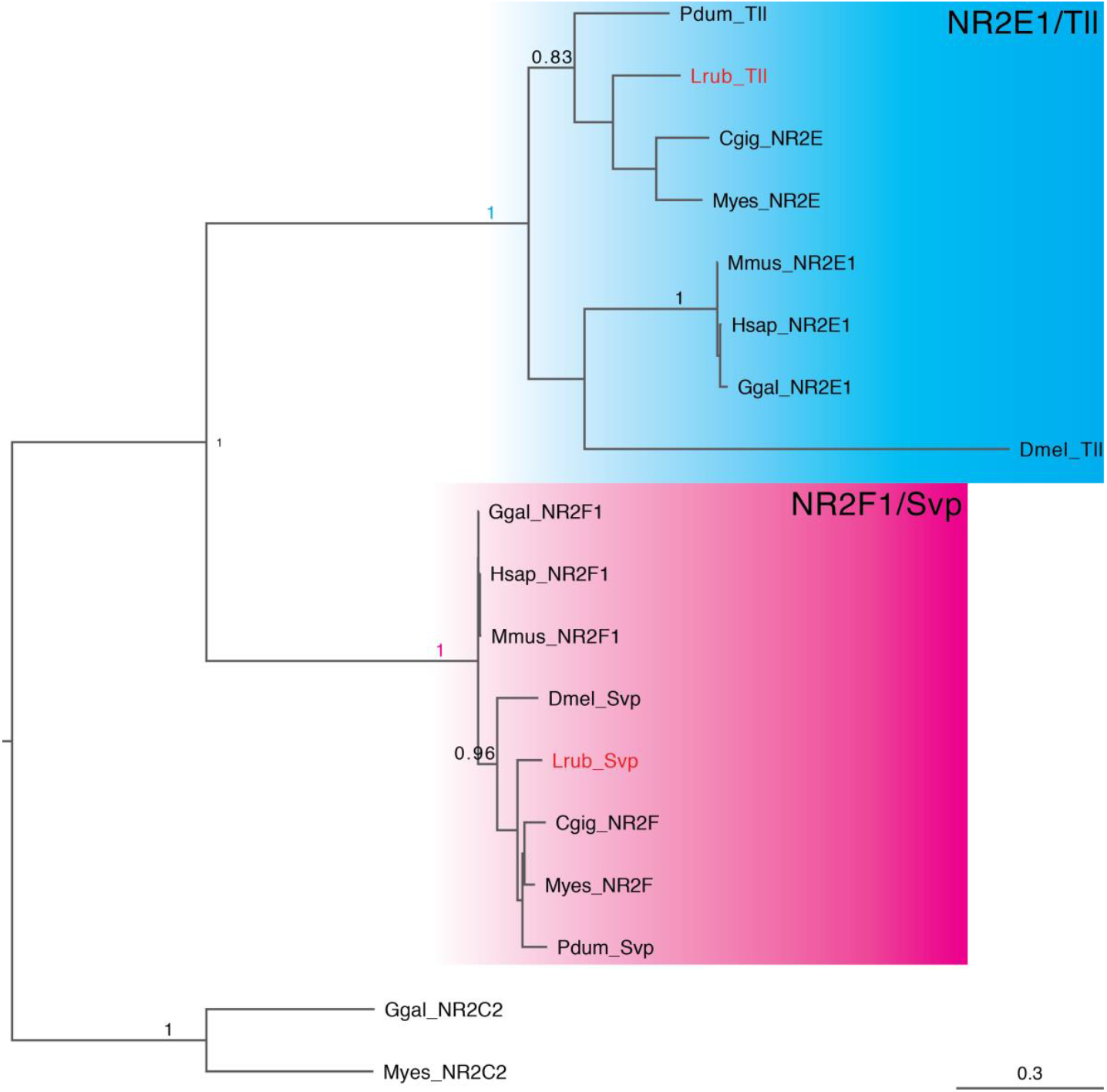
Phylogenetic analysis of nuclear receptor subfamily 2. SH-like support values are shown for the important nodes. Scale bar on the lower right corner shows amino acid substitution rate per site. Sequences from *L. ruber* are marked in red. For abbreviation and source of other sequences see table S1.

**Table S1.**
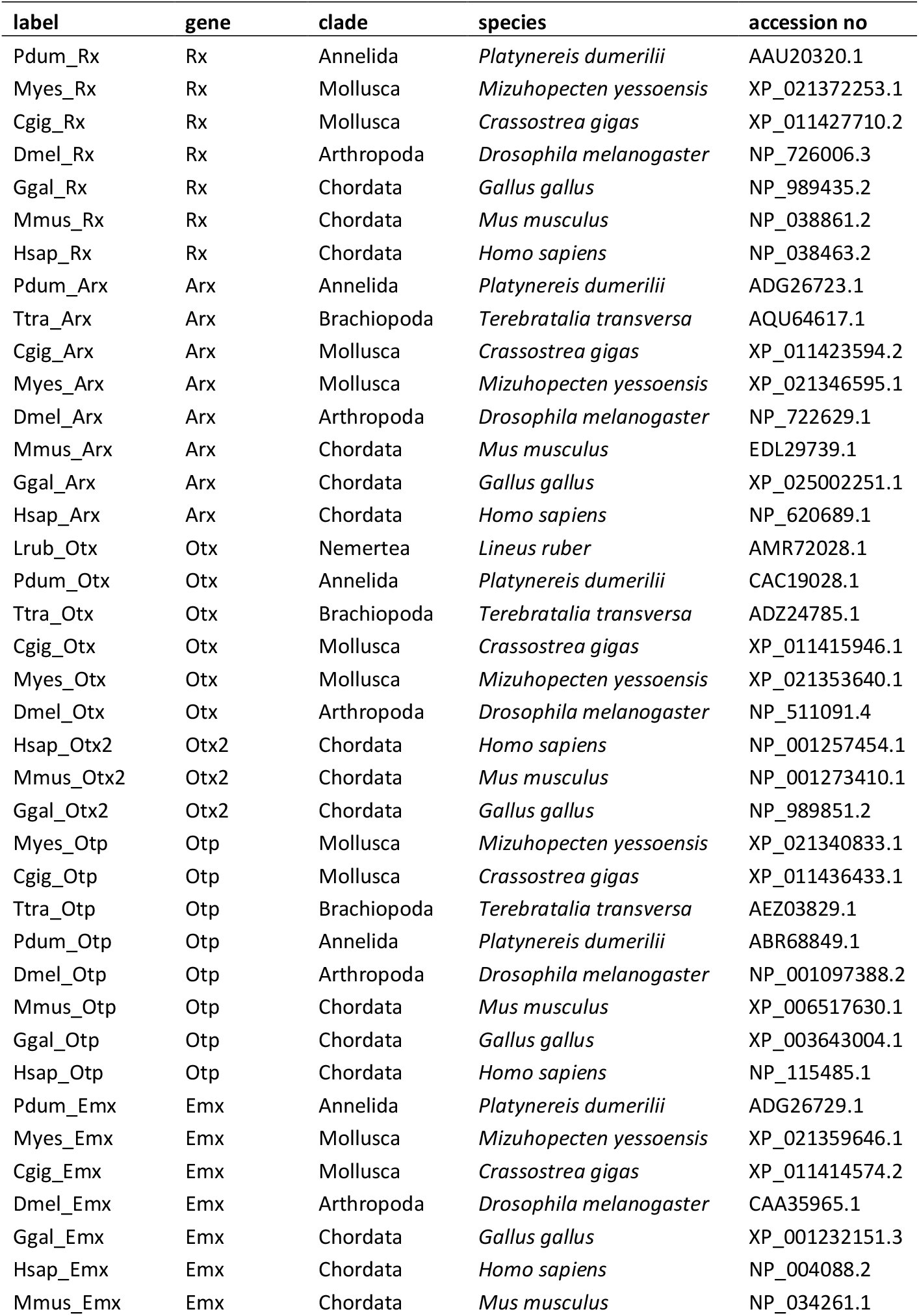
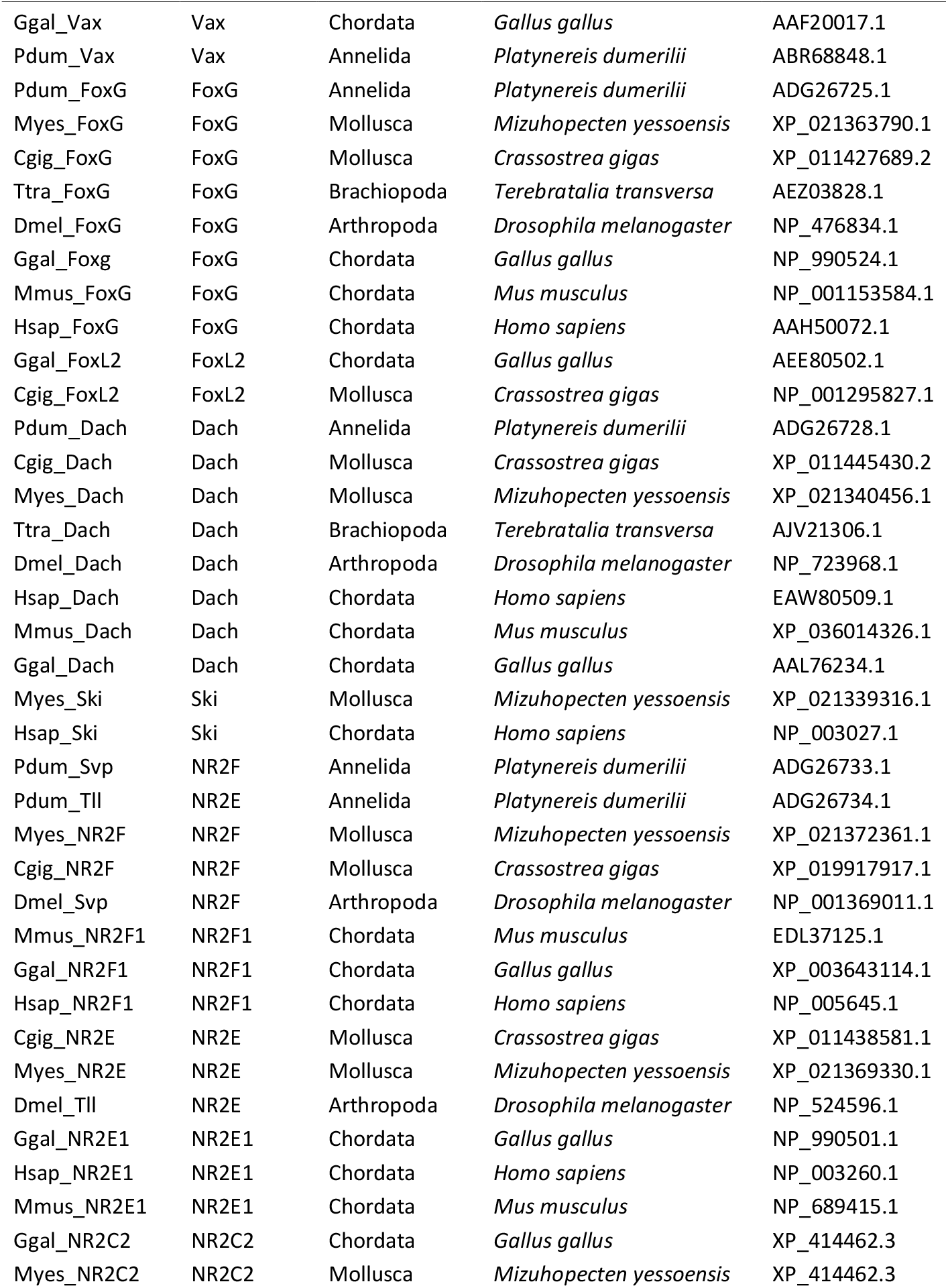
Sequences used in phylogenetic analyses

