## Additional File 1 for "Gene expression in the developing nemertean brain indicates convergent evolution of complex brains in Spiralia"

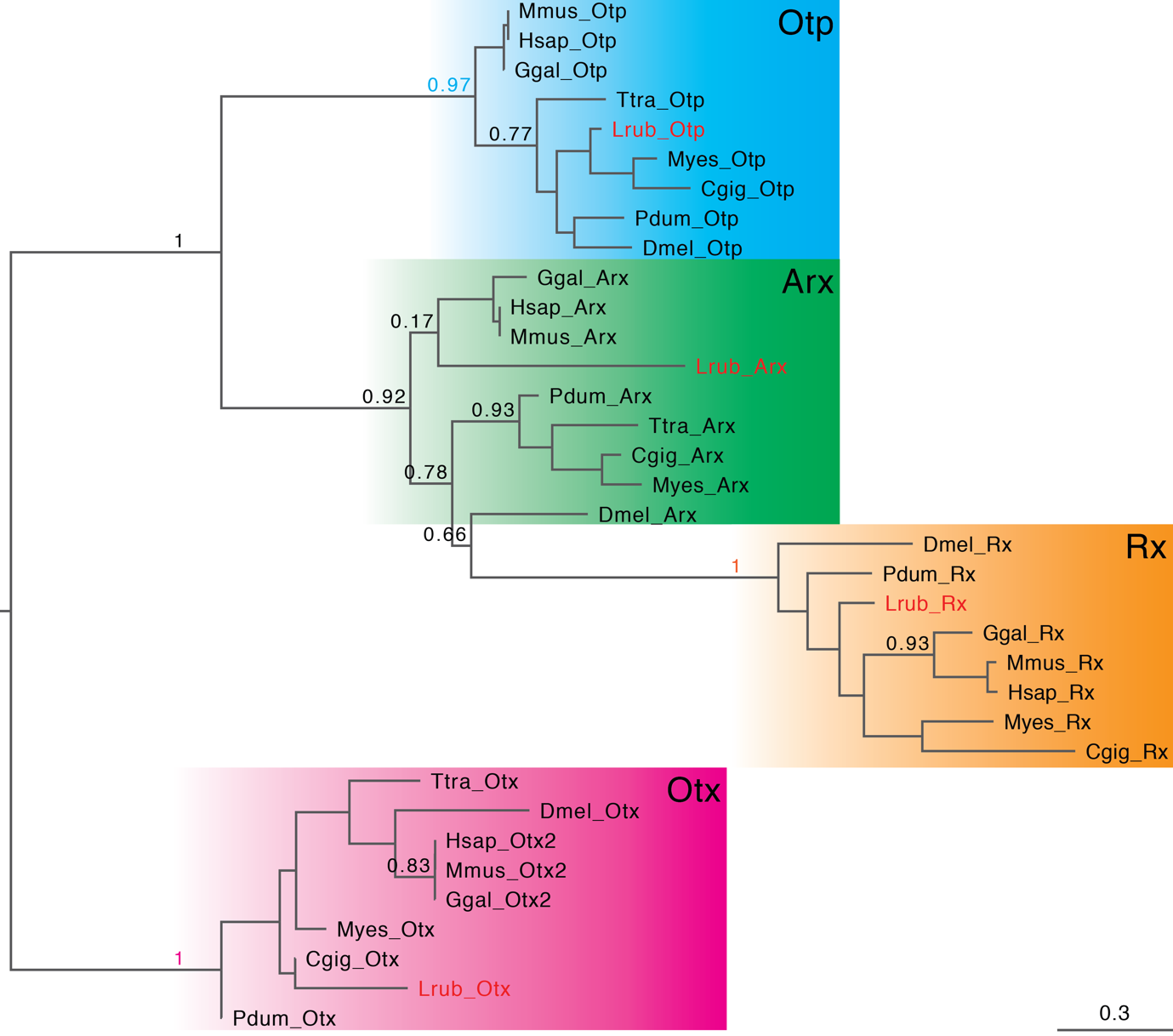

**Fig. S1.** Phylogenetic analysis of PRD-class homeobox transcription factors. SH-like support values are shown for the important nodes. Scale bar on the lower right corner shows amino acid substitution rate per site. Sequences from *L. ruber* are marked in red. For abbreviation and source of other sequences see table S1.

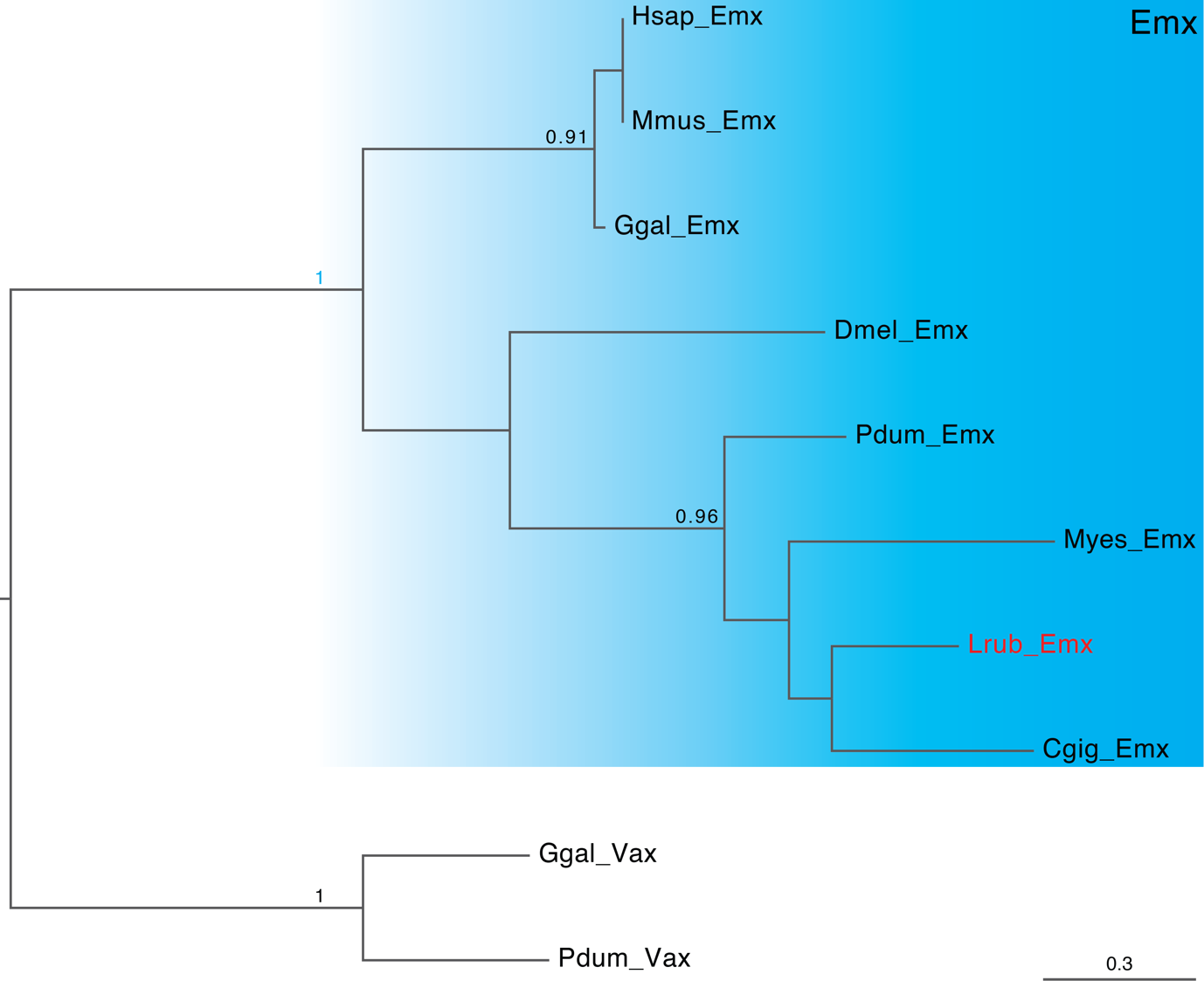

**Fig. S2.** Phylogenetic analysis of Emx sequences. SH-like support values are shown for the important nodes. Scale bar on the lower right corner shows amino acid substitution rate per site. Sequence from *L. ruber* is marked in red. For abbreviation and source of other sequences see table S1.

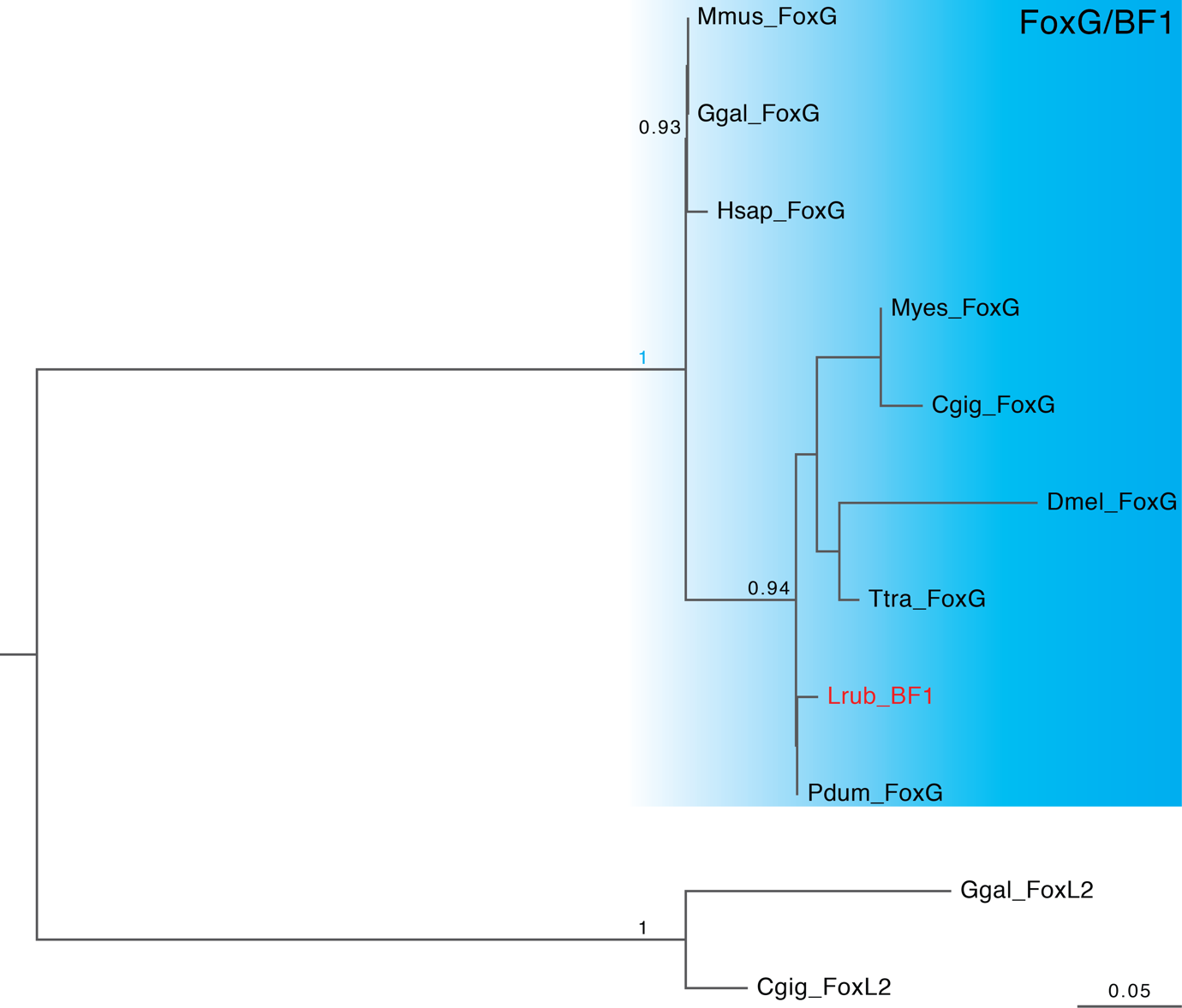

**Fig. S3.** Phylogenetic analysis of Fox sequences. SH-like support values are shown for the important nodes. Scale bar on the lower right corner shows amino acid substitution rate per site. Sequence from *L. ruber* is marked in red. For abbreviation and source of other sequences see table S1.

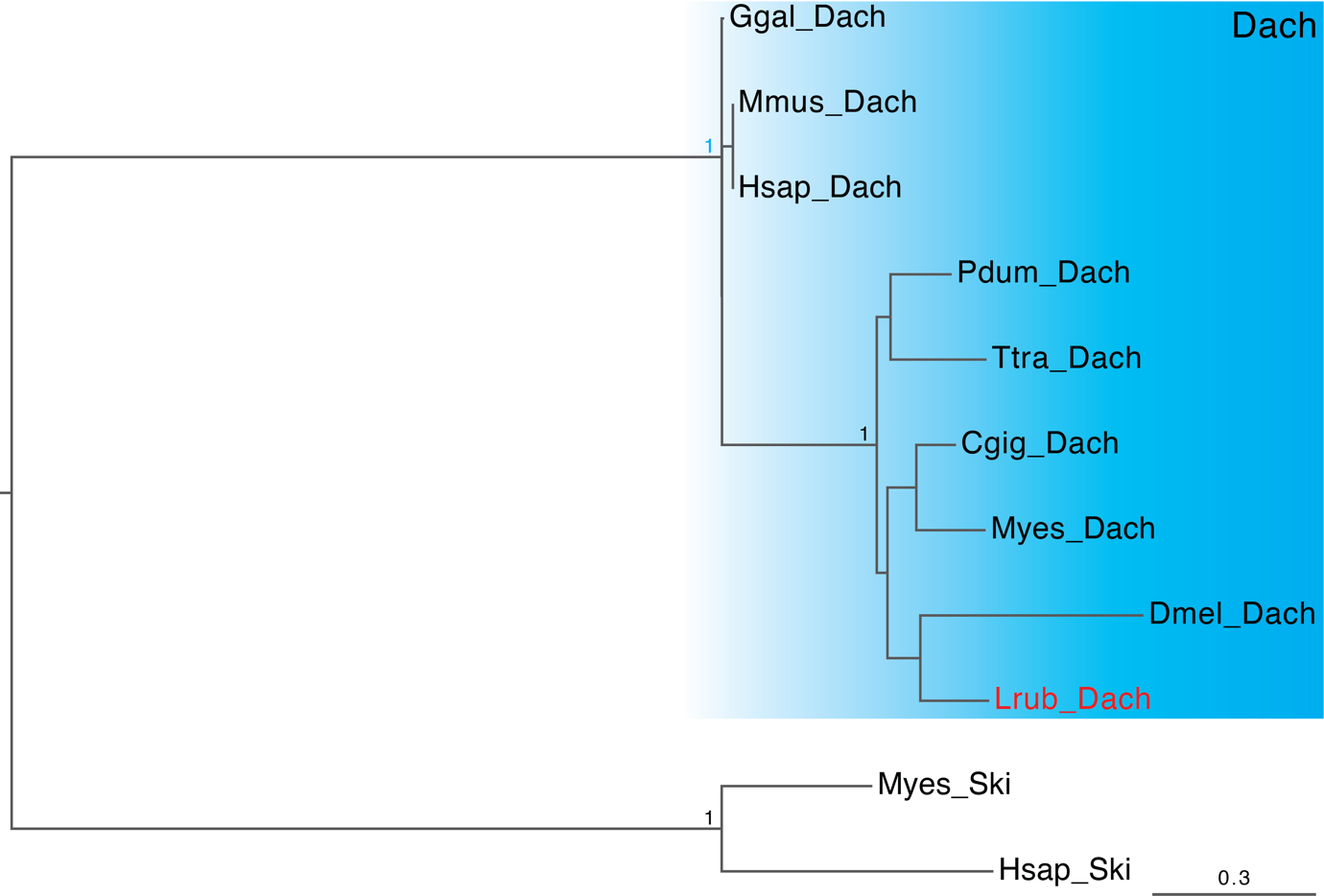

**Fig. S4.** Phylogenetic analysis of Dach sequences. SH-like support values are shown for the important nodes. Scale bar on the lower right corner shows amino acid substitution rate per site. Sequence from *L. ruber* is marked in red. For abbreviation and source of other sequences see table S1.

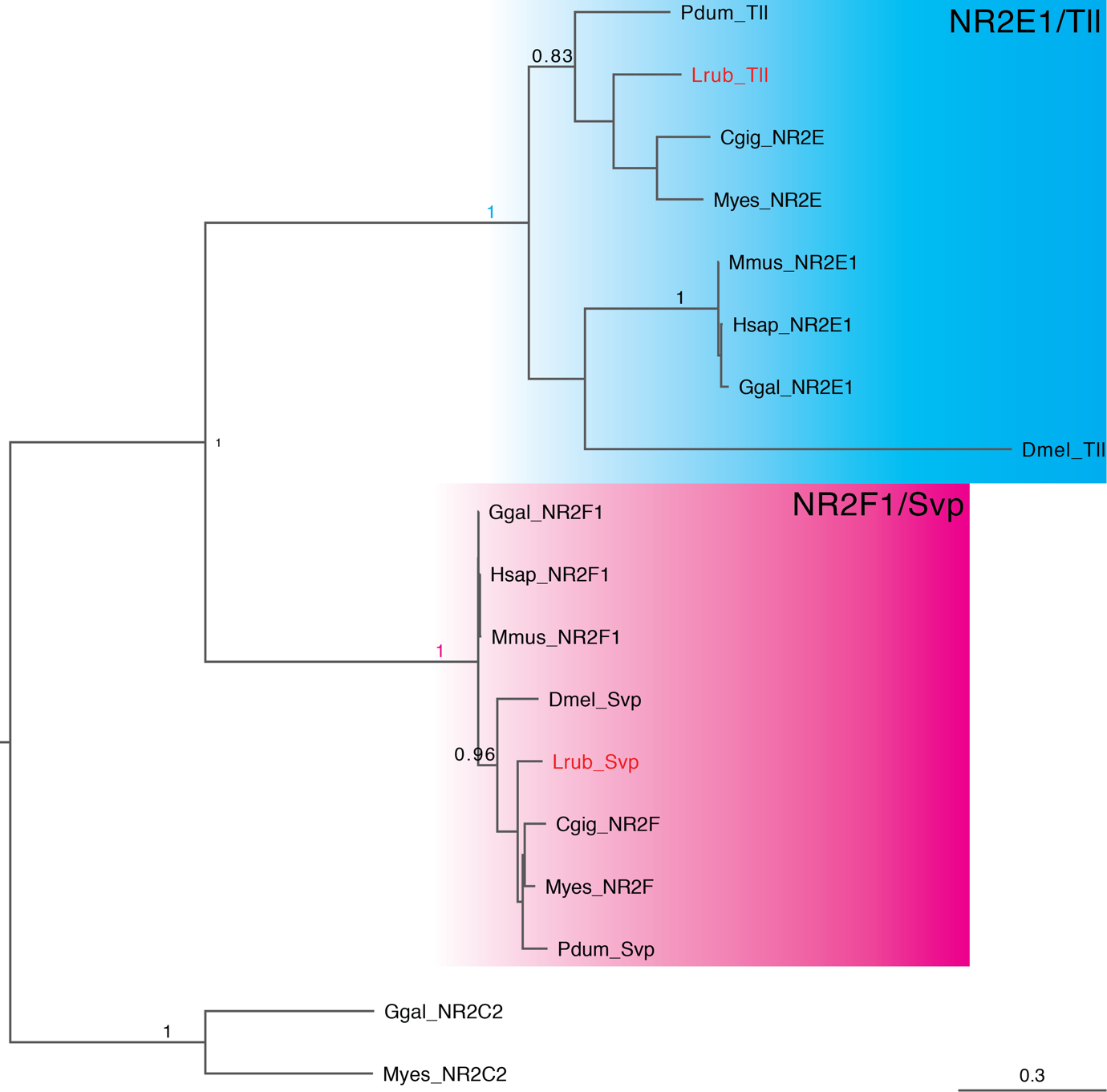

**Fig. S5.** Phylogenetic analysis of nuclear receptor subfamily 2. SH-like support values are shown for the important nodes. Scale bar on the lower right corner shows amino acid substitution rate per site. Sequences from *L. ruber* are marked in red. For abbreviation and source of other sequences see table S1.

**Table S1.** Sequences used in phylogenetic analyses

| **label** | **gene** | **clade** | **species** | **accession no** |
| --- | --- | --- | --- | --- |
| Pdum_Rx | Rx | Annelida | *Platynereis dumerilii* | AAU20320.1 |
| Myes_Rx | Rx | Mollusca | *Mizuhopecten yessoensis* | XP_021372253.1 |
| Cgig_Rx | Rx | Mollusca | *Crassostrea gigas* | XP_011427710.2 |
| Dmel_Rx | Rx | Arthropoda | *Drosophila melanogaster* | NP_726006.3 |
| Ggal_Rx | Rx | Chordata | *Gallus gallus* | NP_989435.2 |
| Mmus_Rx | Rx | Chordata | *Mus musculus* | NP_038861.2 |
| Hsap_Rx | Rx | Chordata | *Homo sapiens* | NP_038463.2 |
| Pdum_Arx | Arx | Annelida | *Platynereis dumerilii* | ADG26723.1 |
| Ttra_Arx | Arx | Brachiopoda | *Terebratalia transversa* | AQU64617.1 |
| Cgig_Arx | Arx | Mollusca | *Crassostrea gigas* | XP_011423594.2 |
| Myes_Arx | Arx | Mollusca | *Mizuhopecten yessoensis* | XP_021346595.1 |
| Dmel_Arx | Arx | Arthropoda | *Drosophila melanogaster* | NP_722629.1 |
| Mmus_Arx | Arx | Chordata | *Mus musculus* | EDL29739.1 |
| Ggal_Arx | Arx | Chordata | *Gallus gallus* | XP_025002251.1 |
| Hsap_Arx | Arx | Chordata | *Homo sapiens* | NP_620689.1 |
| Lrub_Otx | Otx | Nemertea | *Lineus ruber* | AMR72028.1 |
| Pdum_Otx | Otx | Annelida | *Platynereis dumerilii* | CAC19028.1 |
| Ttra_Otx | Otx | Brachiopoda | *Terebratalia transversa* | ADZ24785.1 |
| Cgig_Otx | Otx | Mollusca | *Crassostrea gigas* | XP_011415946.1 |
| Myes_Otx | Otx | Mollusca | *Mizuhopecten yessoensis* | XP_021353640.1 |
| Dmel_Otx | Otx | Arthropoda | *Drosophila melanogaster* | NP_511091.4 |
| Hsap_Otx2 | Otx2 | Chordata | *Homo sapiens* | NP_001257454.1 |
| Mmus_Otx2 | Otx2 | Chordata | *Mus musculus* | NP_001273410.1 |
| Ggal_Otx2 | Otx2 | Chordata | *Gallus gallus* | NP_989851.2 |
| Myes_Otp | Otp | Mollusca | *Mizuhopecten yessoensis* | XP_021340833.1 |
| Cgig_Otp | Otp | Mollusca | *Crassostrea gigas* | XP_011436433.1 |
| Ttra_Otp | Otp | Brachiopoda | *Terebratalia transversa* | AEZ03829.1 |
| Pdum_Otp | Otp | Annelida | *Platynereis dumerilii* | ABR68849.1 |
| Dmel_Otp | Otp | Arthropoda | *Drosophila melanogaster* | NP_001097388.2 |
| Mmus_Otp | Otp | Chordata | *Mus musculus* | XP_006517630.1 |
| Ggal_Otp | Otp | Chordata | *Gallus gallus* | XP_003643004.1 |
| Hsap_Otp | Otp | Chordata | *Homo sapiens* | NP_115485.1 |
| Pdum_Emx | Emx | Annelida | *Platynereis dumerilii* | ADG26729.1 |
| Myes_Emx | Emx | Mollusca | *Mizuhopecten yessoensis* | XP_021359646.1 |
| Cgig_Emx | Emx | Mollusca | *Crassostrea gigas* | XP_011414574.2 |
| Dmel_Emx | Emx | Arthropoda | *Drosophila melanogaster* | CAA35965.1 |
| Ggal_Emx | Emx | Chordata | *Gallus gallus* | XP_001232151.3 |
| Hsap_Emx | Emx | Chordata | *Homo sapiens* | NP_004088.2 |
| Mmus_Emx | Emx | Chordata | *Mus musculus* | NP_034261.1 |

**Table S1.** Continued.

| **label** | **gene** | **clade** | **species** | **accession no** |
| --- | --- | --- | --- | --- |
| Ggal_Vax | Vax | Chordata | *Gallus gallus* | AAF20017.1 |
| Pdum_Vax | Vax | Annelida | *Platynereis dumerilii* | ABR68848.1 |
| Pdum_FoxG | FoxG | Annelida | *Platynereis dumerilii* | ADG26725.1 |
| Myes_FoxG | FoxG | Mollusca | *Mizuhopecten yessoensis* | XP_021363790.1 |
| Cgig_FoxG | FoxG | Mollusca | *Crassostrea gigas* | XP_011427689.2 |
| Ttra_FoxG | FoxG | Brachiopoda | *Terebratalia transversa* | AEZ03828.1 |
| Dmel_FoxG | FoxG | Arthropoda | *Drosophila melanogaster* | NP_476834.1 |
| Ggal_Foxg | FoxG | Chordata | *Gallus gallus* | NP_990524.1 |
| Mmus_FoxG | FoxG | Chordata | *Mus musculus* | NP_001153584.1 |
| Hsap_FoxG | FoxG | Chordata | *Homo sapiens* | AAH50072.1 |
| Ggal_FoxL2 | FoxL2 | Chordata | *Gallus gallus* | AEE80502.1 |
| Cgig_FoxL2 | FoxL2 | Mollusca | *Crassostrea gigas* | NP_001295827.1 |
| Pdum_Dach | Dach | Annelida | *Platynereis dumerilii* | ADG26728.1 |
| Cgig_Dach | Dach | Mollusca | *Crassostrea gigas* | XP_011445430.2 |
| Myes_Dach | Dach | Mollusca | *Mizuhopecten yessoensis* | XP_021340456.1 |
| Ttra_Dach | Dach | Brachiopoda | *Terebratalia transversa* | AJV21306.1 |
| Dmel_Dach | Dach | Arthropoda | *Drosophila melanogaster* | NP_723968.1 |
| Hsap_Dach | Dach | Chordata | *Homo sapiens* | EAW80509.1 |
| Mmus_Dach | Dach | Chordata | *Mus musculus* | XP_036014326.1 |
| Ggal_Dach | Dach | Chordata | *Gallus gallus* | AAL76234.1 |
| Myes_Ski | Ski | Mollusca | *Mizuhopecten yessoensis* | XP_021339316.1 |
| Hsap_Ski | Ski | Chordata | *Homo sapiens* | NP_003027.1 |
| Pdum_Svp | NR2F | Annelida | *Platynereis dumerilii* | ADG26733.1 |
| Pdum_Tll | NR2E | Annelida | *Platynereis dumerilii* | ADG26734.1 |
| Myes_NR2F | NR2F | Mollusca | *Mizuhopecten yessoensis* | XP_021372361.1 |
| Cgig_NR2F | NR2F | Mollusca | *Crassostrea gigas* | XP_019917917.1 |
| Dmel_Svp | NR2F | Arthropoda | *Drosophila melanogaster* | NP_001369011.1 |
| Mmus_NR2F1 | NR2F1 | Chordata | *Mus musculus* | EDL37125.1 |
| Ggal_NR2F1 | NR2F1 | Chordata | *Gallus gallus* | XP_003643114.1 |
| Hsap_NR2F1 | NR2F1 | Chordata | *Homo sapiens* | NP_005645.1 |
| Cgig_NR2E | NR2E | Mollusca | *Crassostrea gigas* | XP_011438581.1 |
| Myes_NR2E | NR2E | Mollusca | *Mizuhopecten yessoensis* | XP_021369330.1 |
| Dmel_Tll | NR2E | Arthropoda | *Drosophila melanogaster* | NP_524596.1 |
| Ggal_NR2E1 | NR2E1 | Chordata | *Gallus gallus* | NP_990501.1 |
| Hsap_NR2E1 | NR2E1 | Chordata | *Homo sapiens* | NP_003260.1 |
| Mmus_NR2E1 | NR2E1 | Chordata | *Mus musculus* | NP_689415.1 |
| Ggal_NR2C2 | NR2C2 | Chordata | *Gallus gallus* | XP_414462.3 |
| Myes_NR2C2 | NR2C2 | Mollusca | *Mizuhopecten yessoensis* | XP_414462.3 |
